# MOCHA: advanced statistical modeling of scATAC-seq data enables functional genomic inference in large human disease cohorts

**DOI:** 10.1101/2023.06.23.544827

**Authors:** Samir Rachid Zaim, Mark-Phillip Pebworth, Imran McGrath, Lauren Okada, Morgan Weiss, Julian Reading, Julie L. Czartoski, Troy R. Torgerson, M. Juliana McElrath, Thomas F. Bumol, Peter J. Skene, Xiao-jun Li

## Abstract

Single-cell assay for transposase-accessible chromatin using sequencing (scATAC-seq) has been increasingly used to study gene regulation. However, major analytical gaps limit its utility in studying gene regulatory programs in complex diseases. We developed MOCHA (Model-based single cell Open CHromatin Analysis) with major advances over existing analysis tools, including: 1) improved identification of sample-specific open chromatin, 2) proper handling of technical drop-out with zero-inflated methods, 3) mitigation of false positives in single cell analysis, 4) identification of alternative transcription-starting-site regulation, and 5) transcription factor–gene network construction from longitudinal scATAC-seq data. These advances provide a robust framework to study gene regulatory programs in human disease. We benchmarked MOCHA with four state-of-the-art tools to demonstrate its advances. We also constructed cross-sectional and longitudinal gene regulatory networks, identifying potential mechanisms of COVID-19 response. MOCHA provides researchers with a robust analytical tool for functional genomic inference from scATAC-seq data.

## Introduction

Single-cell assay for transposase-accessible chromatin using sequencing (scATAC-seq)^1–3^ has become increasingly popular in recent years for studying biological and translational questions around gene regulation and cell identity and has revealed insights on diverse topics such as tumor-related T cell exhaustion^4^, trained immunity in monocytes in patients with COVID-19^5^, regulators of innate immunity in COVID-19^6^, and potentially causal variants for Alzheimer’s disease and Parkinson’s disease^7^. Many sophisticated software tools have been developed for analyzing scATAC-seq data^8^, covering functionalities such as dimensionality reduction and clustering^9–11^, semi-automated cell type annotation^10,11^, identification of open chromatin regions^12–16^, characterization of motif usage and enrichment^17–20^, and inference on gene regulatory networks^11,21,22^. Integrating these developments, recent end-to-end analysis pipelines streamline the analytical process from quality control and cell type annotation to accessibility and motif analysis^9–12,23^. Together these tools have facilitated the extraction of biological insights from scATAC-seq data.

Despite these advances, major analytical gaps in scATAC-seq data analysis limit the construction of robust and reproducible gene regulatory networks to study human disease. First, human disease studies require reliable evaluation of sample- and cell-type-specific open chromatin to capture human genetic heterogeneity and cell type-specific regulatory regions. However, visibility into these forms of heterogeneity is compromised by existing packages^10–13,15^, which usually mix cells across either samples or cell types to compensate for the low coverage of scATAC-seq. Second, scATAC-seq data is intrinsically sparse. Only 5–15% of open chromatin regions are detected in individual cells^9^. Both single-cell and pseudo-bulk ATAC-seq data can contain an excessively high number of regions without accessibility measurements. While zero-inflated (ZI) statistical methods are widely used in analyzing single-cell ribonucleic acid sequencing (scRNA-seq) data^24–27^, such methods are not implemented in popular tools for scATAC-seq data analysis, likely leading to many unreliable results. Third, previous studies have shown that pseudo-replication bias (cell-interdependence) generates many false results in scRNA-seq analysis, if left unaddressed^28,29^. Similarly, any scATAC-seq tools that do not address this issue may generate many false results as well. In longitudinal studies, this pseudo-replication bias is exacerbated as subjects have multiple interdependent samples. To postulate robust gene regulatory networks in human disease, the research community needs a tool to address these challenges.

To this end, we developed a suite of analytical modules for robust functional genomic inference in heterogeneous human disease cohorts, in an open R package called MOCHA (Model-based single cell Open CHromatin Analysis). First, we developed a method for evaluating sample- and cell-type-specific chromatin accessibility in low coverage scATAC-seq. Second, we implemented ZI statistical methods^30–34^ for differential accessibility analysis, co-accessibility analysis, and mixed effects modeling. Third, we aggregated scATAC-seq data per cell type into normalized pseudo-bulk tile-sample accessibility matrices (TSAMs) to capture donor and cell type-centric accessibility. This TSAM directly addresses the issue of cell interdependence (pseudo-replication) and enables modeling cross-sectional and longitudinal gene regulatory landscapes of human disease in large cohorts. We benchmarked MOCHA against state-of-the-art methods in identifying regions of open chromatin, differential accessibility, and co-accessibility. More importantly, we demonstrate MOCHA’s ability to construct gene regulatory networks from both cross-sectional and longitudinal analyses of scATAC-seq data on CD16 monocytes from our COVID-19 cohort^35^. We also demonstrate how to integrate MOCHA with existing tools^11,17,30,36^, and adapt it for custom approaches. In all, we anticipate MOCHA will accelerate the analysis and interpretation of gene regulatory networks using scATAC-seq data.

## Results

### MOCHA overview

We developed MOCHA based on our COVID19 dataset (Methods), which was collected on n=91 peripheral blood mononuclear cell (PBMC) samples of either COVID+ participants (n=18, 10 females and 8 males, 3–5 samples per participant, a total of 69 samples) or uninfected COVID- participants (n=22, 10 females and 12 males, one sample per participant). We obtained high quality scATAC-seq data of 1,311,638 cells from the samples. Unless specified, we mainly used a cross-sectional subset of the COVID19 dataset (denoted as COVID19X, n=39) in the development, including data of the COVID- samples and the first samples of the COVID+ participants during early infection (<16 days post symptom onset (PSO), n=17, 9 females and 8 males).

We designed MOCHA to serve as an analytical framework for sample-centric scATAC-seq data analysis, after quality control assessments and cell type labeling. MOCHA identifies sample- and cell type-specific open chromatin and provides a range of analytical functions for complex scATAC-seq data analysis (Fig. 1, Supplementary Fig. 1):

i. Tiling: MOCHA divides the genome into pre-defined 500 base pair (bp) tiles, which allows head-to-head comparisons on chromatin accessibility across samples and cell types and avoids complex peak-merging procedure on non-aligned summits^9,11^.
ii. Normalization: Since sequencing depth may differ across samples in a large-scale scATAC-seq study (Supplementary Fig. 2a), it is essential to normalize scATAC-seq data prior to meaningful accessibility analysis. MOCHA counts the number of fragments that overlap with individual tiles in individual cells, collects the total and the maximum fragment counts for each tile from all cells of a targeted cell type per sample, and normalizes the fragment counts by the total number of fragments of the cell type per sample to reduce the effects of variations in sequencing depth and cell count (Methods). Compared with other normalization approaches, this approach resulted in the lowest coefficient of variation (CV) distribution on 2230 invariant CCCTC-binding factor (CTCF) sites on the COVID19 dataset (Supplementary Fig. 2b, n=91). As indicated by the low to moderate CV values, this approach also makes it possible to compare normalized accessibility across samples (Supplementary Fig. 2b) and cell types (Supplementary Fig. 2c).
iii. Accessibility evaluation: MOCHA applies logistic regression models (LRMs) to evaluate whether a tile in a given cell type and sample is accessible based on three parameters (Methods): the normalized total fragment count *λ*^(1)^, the normalized maximum fragment count *λ*^(2)^, and a study-specific prefactor *S* to account for differences in data quality between training and user datasets. Since *λ*^(3)^ can only be evaluated on scATAC-seq data, its usage distinguishes MOCHA from peak-calling methods based on pseudo-bulk ATAC-seq data only. Using the full COVID19 dataset (n=91), we generated pseudo-bulk ATAC-seq data from scATAC-seq data of all cells of a targeted cell type, ran MACS2^37^ to identify all accessible regions, and used these labels as the imperfect “ground truth” to train and test the LRMs. More specifically, we used natural killer (NK) cells (n=179,836) for training, which had 750 million total fragments, or 15x the recommended coverage for MACS2. On this dataset, MACS2 identified 1.15 million tiles as accessible. Using these tiles as positives and other fragment-containing tiles as negatives, we developed cell count-specific LRMs with varying coefficients and thresholds to account for variations in cell count across samples (Supplementary Fig. 3a-b). MOCHA applies smoothing and interpolation to find the proper coefficients and thresholds for individual datasets. We validated the LRMs (Supplementary Fig. 3c-d) using data of CD14 monocytes (n=135,949), naive B cells (n = 60,595), CD16 monocytes (n=28,525), NK CD56 bright cells (n=14,692), and conventional dendritic cells (cDCs, n=9,915). We used sensitivity, specificity, area under the receiver operating characteristic (ROC) curve (AUC), and Youden’s J statistic to quantify the performance. Overall MOCHA had a good performance even at low cell counts. For example, MOCHA achieved an AUC ranging from 0.693 (CD14 monocytes), 0.703 (CD16 monocytes), to 0.741 (NK CD56 bright cells) with only 50 cells.
iv. Tile-sample accessibility matrix (TSAM): MOCHA first uses the LRMs to identify accessible tiles from cells of a targeted cell type in individual samples and then keeps only tiles that are common to at least a user-defined fraction threshold of samples. Afterwards, MOCHA generates a TSAM for the cell type with rows being the accessible tiles, columns the samples, and elements the corresponding *λ*^(1)^ values. A total of 215,649 accessible tiles were identified on CD16 monocytes with a fraction threshold of 20% (Supplementary Fig. 2d) across either COVID+ or COVID- samples in the COVID19X dataset (n=39). About 25% elements in the obtained TSAM were zero (Supplementary Fig. 2e,f), reflecting the sparsity of scATAC-seq data even after pseudo-bulking and indicating the necessity of applying ZI statistical methods for downstream analysis.
v. Differential accessibility analysis (DAA): Similar to other methods, MOCHA first filters out noisy tiles. Tiles are excluded if either 1) the median *log2*(*λ*^(1)^ + 1) value (across all samples) is lower than a user-defined threshold or 2) their difference (between two sample groups) in percentage of zeros is less than a user-defined threshold. Unlike other methods, MOCHA includes functions to heuristically define these thresholds. For the COVID19X dataset (n=39), we noticed that the *log2*(*λ*^(1)^ + 1) values in the TSAM of CD16 monocytes followed a bi-modal model and thus chose a value of 12 near the higher mode as the median accessibility threshold (Supplementary Fig. 2g). Additionally, we observed a 25% difference in fragment counts between COVID+ and COVID- samples (Supplementary Fig. 2a), so we set a 50% threshold for the difference in the percentage of zeros to control for technical differences. MOCHA then applies a two-part Wilcoxon test ^31,32^ to identify differential accessibility tiles (DATs) within the cell type between the two sample groups (Methods). DATs have either a large fold change (FC) in accessibility (Supplementary Fig. 2h) and/or large difference in percentage of zeros (Supplementary Fig. 2i). For comparison, ArchR^11^ uses the standard Wilcoxon test along with a post-test log2(FC) cutoff to identify differential regions on bias-matched cell populations, while Signac^10^ constructs LRMs and prioritizes differential regions based on FC.
vi. Co-accessibility analysis (CAA): MOCHA applies the ZI Spearman correlation^33^ to evaluate two types of co-accessibility between tiles in TSAMs (Methods). The inter-cell type co-accessibility is evaluated across cell types where data from different samples are stacked. The inter-sample co-accessibility is evaluated within a targeted cell type but across different samples. Both types are important to infer potential gene regulatory networks, one for understanding differences between cell types and the other for understanding differences between sample groups.

**Figure 1.**
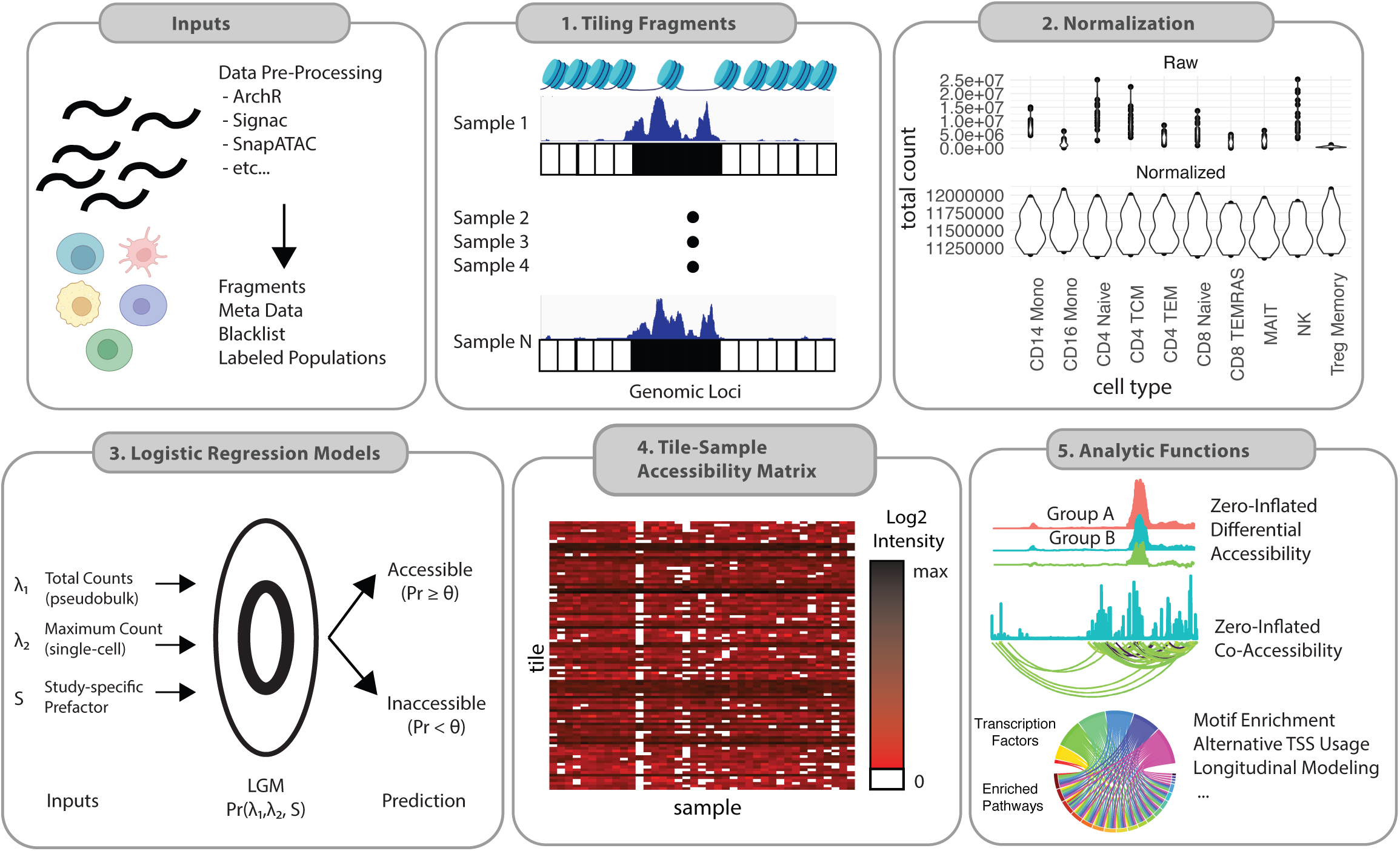
General workflow of MOCHA. Schematic representation of the core functionalities in MOCHA, starting from scATAC input data (fragments, black list, cell type labels, and sample metadata). Using these data, MOCHA generates fragment counts for every 500 bp tiles (1), normalizes the count data (2), and leverages single-cell and pseudo-bulk information to identify open tiles in a cell type- and sample-specific manner (3). It then generates population-level open chromatin matrices for each cell type (4), which is the starting point for downstream analytical functions (5). MOCHA includes improvements to differential accessibility analysis, co-accessibility analysis, and longitudinal modeling. It also provides functions for identifying alternatively regulated transcription starting sites, motif enrichment, and dimensionality reduction. Figures were generated using Adobe Illustrator and BioRender.

In addition, MOCHA has functions for dimensionality reduction, motif enrichment, analysis for alternative transcription starting site (TSS) regulation, and longitudinal modeling. TSAMs can also be passed to some existing scATAC-seq tools such as ArchR and chromVAR and other bioinformatics tools such as Monocle3 for further analysis. Furthermore, users can easily leverage information from TSAMs to conduct their own interrogations of scATAC-seq data.

### MOCHA reliably detects sample-specific chromatin accessibility

A crucial component of scATAC-seq data analysis is to reliably detect which chromatin regions are accessible. We benchmarked MOCHA against the popular tools MACS2^37^ and HOMER^38^. The former is also implemented in ArchR^11^, Signac^10^ and SnapATAC^9^. We compared these tools using three scATAC-seq datasets with different data quality and sequencing depth (Methods, Supplementary Fig. 4a): i) COVID19X (n=39, Fig. 2) or COVID19 (n=91, Supplementary Fig. 4); ii) HealthyDonor, a dataset of 18 PBMC samples of n=4 healthy donors ^39^; and iii) Hematopoiesis, an assembled dataset of hematopoietic cells from 49 samples of diverse data quality^11^, which was treated as a single sample sample in this study. Three representative cell types with moderate to high cell counts were selected from each of the three datasets for the comparison, with cell count per sample ranging from 227 to 743 (COVID19X, median), 163 to 784 (HealthyDonor, median), and 1175 to 27463 (Hematopoiesis), respectively (Fig. 2a, Supplementary Fig. 4b).

**Figure 2.**
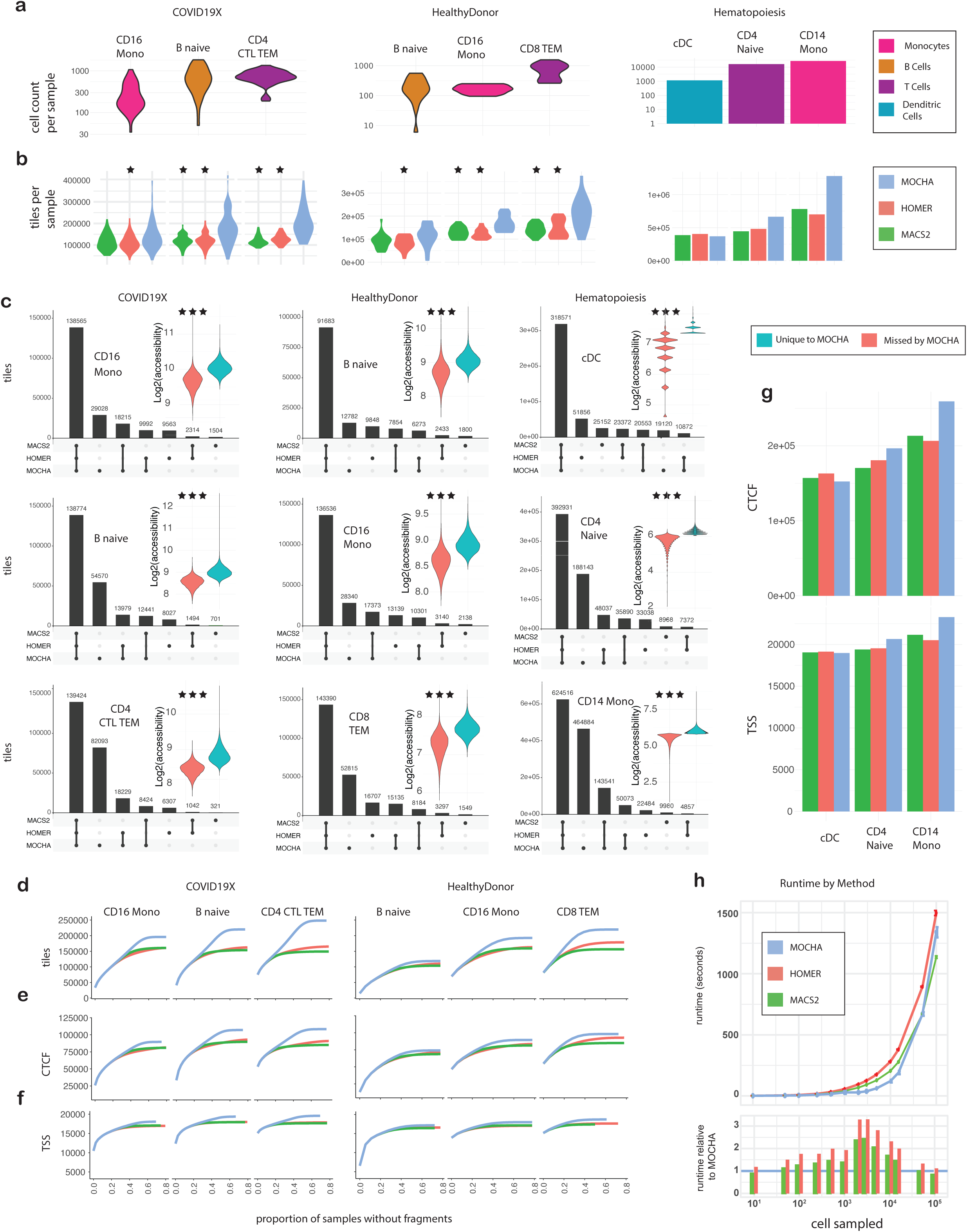
Benchmarking MOCHA with MACS2 and HOMER on open chromatin identification. **a**, Cell counts per sample in three representative cell types from each of three scATAC-seq datasets. The same three cell types in the three corresponding datasets (Methods) were used in this analysis, including COVID19X (n=39), HealthyDonor (n=18, middle panel), and Hematopoiesis (treated as n=1, right panel). **b**, The number of open tiles per sample as identified by MOCHA (light blue), MACS2 (green), or HOMER (red). The same colors are used in **d-h**. **b**,**c**, The Wilcoxon rank sum test was used to compare results by MOCHA with those by MACS2 and/or HOMER. Significantly higher MOCHA values are indicated with * (0.01 < *P* < 0.05) or *** (*P* < 0.001). **c**, UpSet plot showing overlaps between open tiles identified by MOCHA, MACS2, or HOMER. Only tiles common to at least 20% of samples were kept for the COVID19X and HealthyDonor datasets while all identified tiles were kept for the Hematopoiesis dataset. Insert: Violin plot of signals (i.e., log_2_(normalized fragment count+1)) of tiles missed by MOCHA (i.e., those identified by MACS2 and/or HOMER but not by MOCHA, left panel) and signals of tiles unique to MOCHA (i.e., those identified only by MOCHA, right panel). **d-f**, The cumulative number of detected tiles (**d**), tiles overlapping with CCCTC-binding factor (CTCF) sites (**e**), or tiles overlapping with transcription starting sites (TSSs, **f**) as a function of the maximum fraction of samples allowed to have no fragments in the COVID19X (left panel) or HealthyControl (right panel) datasets. **g**, The number of detected CTCF sites (top panel) or TSSs (bottom panel) in the Hematopoiesis dataset. **h,** The actual (top panel) and the relative (with respect to MOCHA, bottom panel) runtime required to identify open chromatin from single cell data as a function of the number of downsampled cells. The blue horizontal line at 1 in the bottom panel marks the MOCHA runtime. CD16 Mono: CD16 monocytes; B Naive: naive B cells; CD4 CTL TEM: CD4^+^ cytotoxic T lymphocytes and CD4^+^ effector memory T cells; CD8 TEM: CD8^+^ effector memory T cells; cDC: conventional dendritic cells; CD4 Naive: naive CD4^+^ T cells; CD14 Mono: CD14 monocytes. Source data are provided in Source Data Fig. 2-1 and 2-2. Figures were generated using Adobe Illustrator.

To benchmark performance on sample-specific accessibility, we compared the number of open regions detected in individual samples (Fig. 2b). On COVID19X, MOCHA detected a median of 129k–195k open tiles per sample, which was 19–64% higher (significantly with *P* < 0.05 in ⅚ cases) than the corresponding numbers by MACS2 or HOMER. Similarly on HealthyDonor, MOCHA detected a median of 117k–216k open tiles per sample, a 35–59% increase (significantly with *P* < 0.05 in ⅚ cases) over the corresponding numbers by MACS2 or HOMER. On Hematopoiesis, MOCHA detected 370k open tiles in cDCs, 665k in naive CD4^+^ T cells, and 1.28m in CD14 monocytes, which were <7% lower, >43% higher, and >70% higher, respectively, than the corresponding numbers by MACS2 or HOMER. Thus MOCHA was more sensitive than MACS2 and HOMER in detecting open chromatin regions in individual samples in almost all cases.

To assess the consistency between open tiles detected by the three tools, we generated TSAMs with a fraction threshold of 20% based on tiles detected by each tool and compared the corresponding tiles in TSAMs (Fig. 2c). Most tiles were detected by all three tools. As described above, MOCHA detected more tiles than MACS2 and HOMER in almost all cases. Among all tiles detected by MACS2 and/or HOMER, 95–97% were also detected by MOCHA in COVID19X, 92–93% in HealthyDonor, and 84–97% in Hematopoiesis. Tiles detected only by MOCHA contained on average more fragments than tiles missed by MOCHA (Fig. 2c, inserts; *P* < 0.001). Thus MOCHA not only captured the majority of tiles detected by MACS2 and HOMER but also added extra tiles of better signals than those by MACS2 or HOMER.

To further elucidate differences among tiles detected by the three tools, we calculated the percentage of zeros among all samples for each tile in each TSAM and obtained the corresponding cumulant distributions (Fig. 2d). While all three tools agreed on open tiles common to most samples, MOCHA detected more sample-specific tiles than MACS2 and HOMER. To test whether these additional tiles potentially contained biological information, we generated the corresponding cumulative distributions on tiles mapped to CTCF sites or TSSs and observed similar patterns (Fig. 2e,f). This suggests that the extra tiles detected by MOCHA may carry important biological information. MOCHA also detected similar or more open CTCF sites and open TSSs in Hematopoiesis compared to MACS2 and HOMER (Fig. 2g).

Calling peaks on pooled cells of interest is a common practice in scATAC-seq data analysis^9–11^. To compare MOCHA, MACS2 and HOMER on this approach, we pooled cells of the three cell types in the three datasets, randomly downsampled the cells to a series of predetermined cell counts, and applied the three tools to detect accessible tiles (Supplementary Fig. 4c). MOCHA consistently detected more tiles, more CTCF sites, and more TSSs than MACS2 and HOMER in almost all cases.

Finally, we benchmarked the runtime for each tool on a cloud computing environment. MOCHA was consistently faster than HOMER in all tested cases and MACS2 in all practical cases (Fig. 2h).

### MOCHA implements zero-inflated differential accessibility and co-accessibility analyses

We evaluated MOCHA’s ZI modules against existing state-of-the-art tools for DAA and CAA. We first benchmarked MOCHA with ArchR and Signac on DAA. To ensure a head-to-head comparison, we applied each method to identify DATs between COVID+ (n=17) and COVID- (n=22) participants from the 215,649 tiles in the TSAM of CD16 monocytes in the COVID19X dataset (Methods). MOCHA identified 6211 DATs (false discovery rate (FDR) < 0.2, Fig. 3a, Supplementary Fig. 5a). In comparison, ArchR and Signac detected 6009 and 1266 DATs, respectively (Fig. 3b). While 28% of DATs by MOCHA were in gene promoter regions, the corresponding percentage was 17% for ArchR and 18% for Signac. As a result, MOCHA, ArchR, and Signac identified 1811, 1006, and 228 genes, respectively, with DATs in their promoter regions. These genes were enriched (adjusted *P* < 0.05) in 27, 1, and 1 Reactome pathways^40^ (Fig. 3c), respectively, illustrating a striking distinction by MOCHA. The same trend was also observed for other pathway databases (Supplementary Fig. 5b). Among the 27 Reactome pathways revealed by MOCHA (Fig. 3d), toll-like receptors (TLRs), myeloid differentiation primary response 88 (MyD88), interleukins, and nuclear factor kappa B (NF-κB) pathways all play important roles in innate immune response to viral infection^41^, consistent with the expected functions of CD16 monocytes. Thus DATs by MOCHA revealed more biological insights than those by ArchR or Signac.

**Figure 3.**
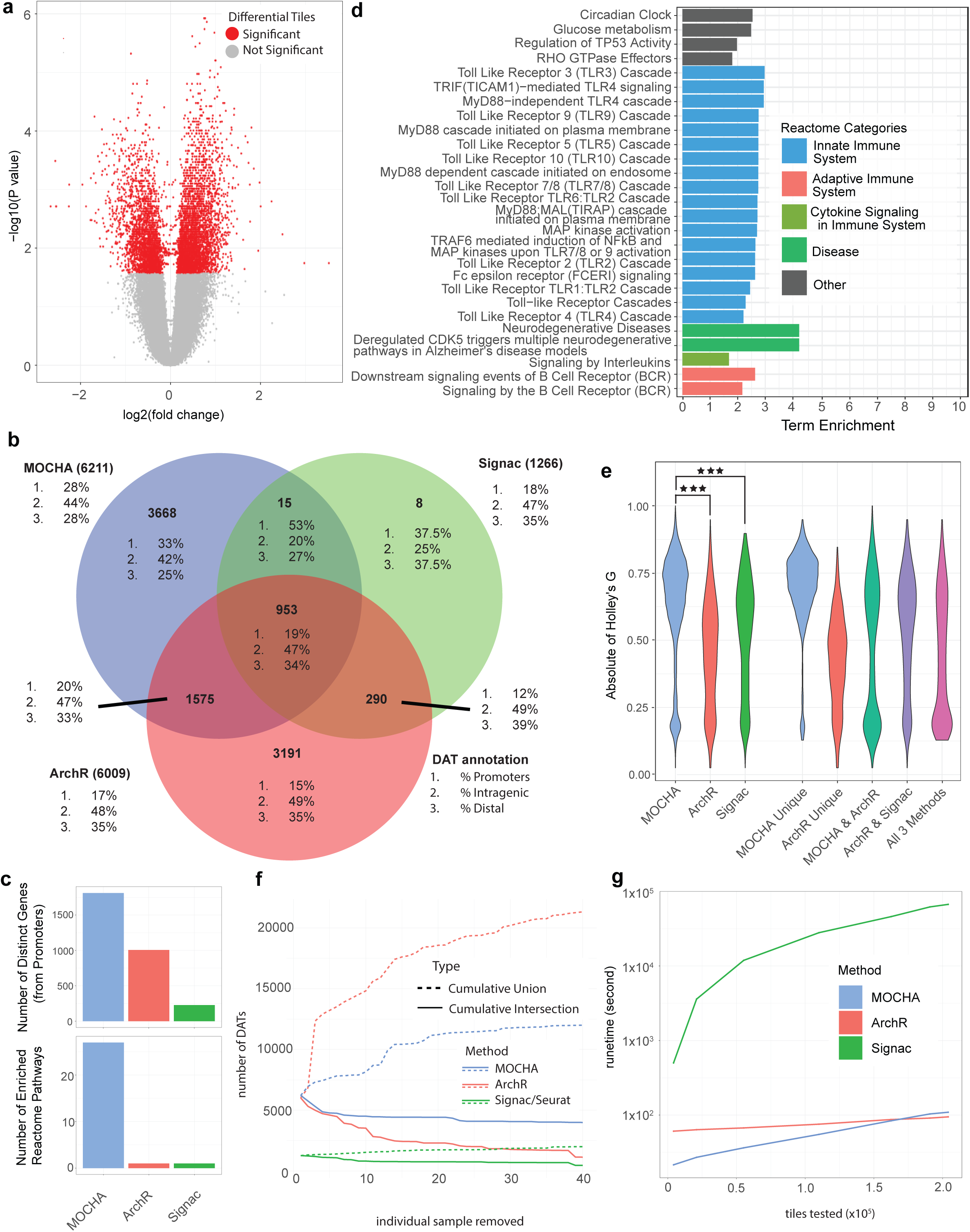
Benchmarking MOCHA with ArchR and Signac on differential accessibility analysis. **a,** MOCHA’s differential accessible tiles (DATs) in CD16 monocytes between COVID+ samples during early infection (n=17) and COVID- samples (n=22) in the COVID19X dataset. The volcano plot illustrates the log2(FC) on the x-axis against the -log10(*P* value) on the y-axis, where FC represents fold change in accessibility. The log2(FC) was estimated using the Hodges-Lehmann estimator^82^. The *P* value was calculated based on the two-part Wilcoxon test^32^. DATs with a false discovery rate (FDR) < 0.2 were considered as significant. **b,** Venn Diagram of MOCHA, Signac, and ArchR’s DATs. The percentage of 1) promoters, 2) intragenic tiles, and 3) distal tiles are shown for each method and each Venn diagram subset. **c,** The number of (top) genes with differential promoters and (bottom) enriched Reactome pathways for each method are depicted using barplots. **d,** Reactome Pathway enrichment results based genes with differential promoter tiles. Pathway categories are annotated using Reactome’s pathway hierarchy. **e,** Violin plot of Holley’s |G| from 1000 bootstrapped samples, each containing 50 randomly selected DATs from each category. Categories with <50 DATs were not tested. ***, *P* < 0.001 (Wilcoxon rank sum test). **f,** Leave-one-sample perturbation analysis to test the robustness of each method in differential accessibility analysis. New sets of DATs were calculated iteratively after removing each sample once. The robustness was assessed by the number of total (solid line) and conserved (dotted line) DATs detected across perturbations. **g,** Each method’s runtime (in seconds) as a function of the number of tested tiles. Source data are provided in Source Data Fig. 3-S5. Figures were generated using Adobe Illustrator.

To quantify the DAT accuracy for each method, we evaluated its efficiency in separating COVID19+ and COVID19- samples. We randomly selected 50 DATs, performed k-mean (k=2) clustering, calculated the absolute value of the G index (|G|) of agreement ^42^, and repeated the process 1000 times. DATs by MOCHA were significantly better in separating COVID19+ and COVID19- samples than those by ArchR (*P* < 0.001) or Signac (*P* < 0.001, Fig. 3e).

Biologically meaningful DATs should be robust against minor changes in the sample set. Starting with DATs from the full sample set (n=39), we iteratively removed one sample at a time and recalculated DATs from the reduced sample set (n=38). We repeated this process until each sample was removed once. From there, we collected the set of conserved (e.g., cumulative intersection) and inflated (e.g., cumulative additional) DATs across all iterations. Ultimately, 3990 (64%) of the 6211 DATs by MOCHA were conserved (Fig. 3f), which was 1.8–3.4 times higher than the corresponding rate of ArchR (1136/6009, 20%) or Signac (458/1264, 36%). MOCHA had 5782 (93%) additional DATs, which was 2.7 times lower than the corresponding rate by ArchR (15310/6009, 255%) and 1.6 times higher than that by Signac (735/1264, 58%). MOCHA was more robust than ArchR and had a split performance in comparison with Signac in detecting DATs regardless of sample set. However, MOCHA had 3990 conserved DATs, 8.7 times higher than those of Signac (458). MOCHA provided a better balance between sensitivity and robustness in detecting DATs compared to ArchR and Signac.

When benchmarking runtime, we observed that MOCHA and ArchR took 1.8 and 1.6 minutes, respectively, to evaluate approximately 200,000 tiles, while Signac took 18.6 hours (Fig. 3g). MOCHA was 23–614x faster than Signac and 0.86–2.8x as fast as ArchR.

Next, we compared the ZI and the standard Spearman correlations across cell types and samples in the COVID19X dataset based on *log_2_*(*λ*^(1)^ + 1) in TSAMs. For the inter-cell type co-accessibility, we used known promoter-enhancer interactions in naive CD4^+^ and CD8^+^ T cells^43^ to define possibly interacting tile pairs (1.21 million) while randomly selecting 100k tile pairs as a negative background for comparison (Methods). Both correlation approaches largely generated similar results (Supplementary Fig. 6a, Spearman correlation = 0.687, *P* < 2.2×10^-16^), but disagreements were also observed (Supplementary Fig. 6b). The ZI correlation better distinguished promoter-enhancer pairs from the random pairs than the standard correlation (Kolmogorov–Smirnov (KS) test statistic = 0.26 vs. 0.13), and identified 1000x more promoter-enhancers pairs (15,988 vs. 149, FDR < 0.1, Supplementary Fig. 6c-d). For the inter-sample co-accessibility, we first collected a subset of tiles in the TSAM of CD16 monocytes that roughly located in the first million bp of chromosome 4 (chr4:121500-1130999) and then used both correlation approaches to calculate the inter-sample correlations between all pairs (about 34.5k) of these tiles. While both correlation approaches generally agreed with each other with a rank correlation of 0.69 (*P* < 0.001, Supplementary Fig. 6e), 9550/34,596 (28%) of the tested correlations switched sign between the two approaches and 2087/34,596 (6%) of them differed in value by >0.25 (Supplementary Fig. 6f). Thus properly accounting for zero-inflation is essential for reliable CAA in sparse scATAC-seq data.

### Networks of alternatively regulated genes in early SARS-CoV-2 infection

To demonstrate how improvements in MOCHA can be leveraged for constructing gene regulatory networks, we investigated possible alternative TSS regulation by CD16 monocytes during early SARS-CoV-2 infection (Methods), using the COVID19X dataset. We observed two types of alternatively regulated genes (ARGs, Fig. 4a): A total of 278 genes had at least one TSS showing differential accessibility (FDR < 0.2) between COVID19+ and COVID19- samples, while other open TSSs had no change (Type I, Fig. 4b). Five genes (ATP1A1, UBAP2L, YWHAZ, CAPN1, and ARHGAP9) had at least two differential TSSs changing in the opposite directions (Type II, Fig. 4c). Interestingly, all Type II ARGs were previously associated with COVID-19 and two (ATP1A1 and CAPN1) have been proposed as therapeutic targets for COVID-19 (Supplementary Table 1). Pathway enrichment analysis^44^ on all Type I and Type II ARGs revealed that they were enriched in pathways of the innate immune response to infection, including MyD88 and TLR responses (Fig. 4d).

**Figure 4.**
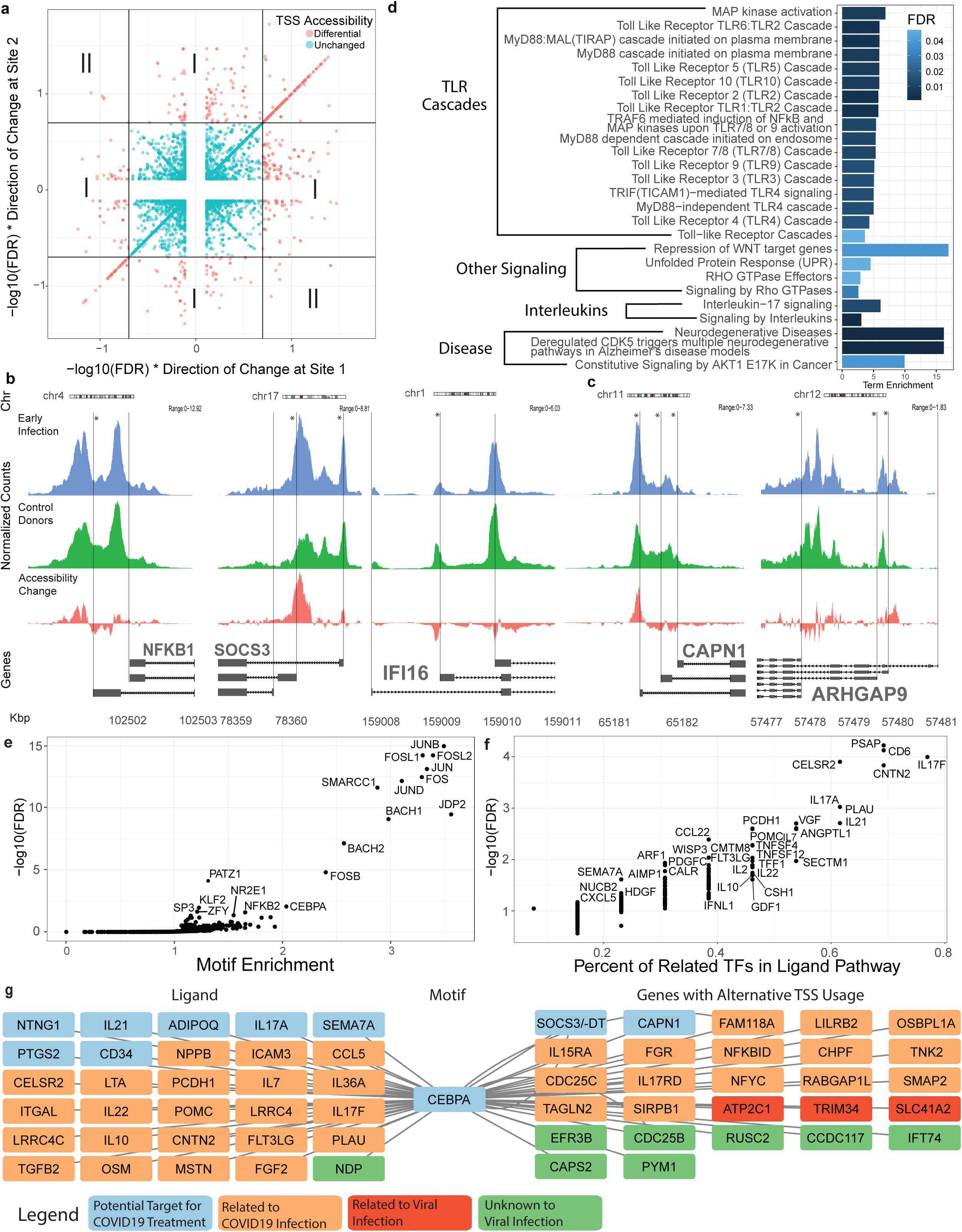
Regulatory network construction on alternative transcription starting sites in CD16 monocytes during early COVID-19 infection. **a,** Scatter plot of differential accessibility at potential alternative transcription starting sites (TSSs). False discovery rate (FDR) and fold change (FC) were evaluated on chromatin accessibility in CD16 monocytes between COVID+ samples during early infection (n=17) and COVID- samples (n=22) in the COVID19X dataset. All pairwise combinations of -log10(FDR)*sign(log2(FC)) are shown. TSS pairs were categorized as type I if only one TSS was significantly differential (FDR < 0.2), or type II if both were significantly differential but in the opposite directions. Pairs of TSSs that were significantly differential in the same direction were not considered. **b-c,** Coverage tracks illustrating type I (**b**) or type II (**c**) alternative TSS regulation around exemplar genes (* denotes a significant differential accessibility tile (DAT), FDR < 0.2). **d,** Reactome pathway enrichment for genes with alternatively regulated TSSs (both type I and II). Pathway annotation was based on Reactome’s hierarchical database. **e,** Motif enrichment using DATs involved in alternatively regulated TSSs and their co-accessible tiles (within ±1M bp, zero-inflated correlation > 0.5). **f,** NicheNet-based ligand-motif set enrichment analysis (LMSEA) on motifs with FDR < 0.01. **g,** A network centered around CEBPA that was constructed using significant ligands, motifs, and genes with alternatively regulated TSS sites. Ligand-motif links represent NicheNet-based associations. Motif-gene links represent motif presence in either an alternative TSS tile, or tiles correlated to an alternative TSS. Source data are provided in Source Data Fig. 4. Figures were generated using Adobe Illustrator.

To understand the upstream signaling mechanisms regulating ARGs, we first applied MOCHA to identify tiles that were co-accessible (inter-sample) with the corresponding DATs. Motif enrichment analysis on DATs of ARGs and their co-accessible tiles identified 13 enriched motifs, including activator protein-1 (AP-1) family motifs, PATZ1, and CEBPA (adjusted P < 0.05, Fig. 4e). Next, we carried out ligand-motif set enrichment analysis (LMSEA, Methods) based on a priori ligand-motif (transcription factor, TF) interactions in NicheNet^45^. We identified 122 significantly enriched ligands (adjusted P < 0.05, Fig. 4f), many of which, including IL17, IL21 and PLAU, have already been implicated in COVID-19 (Source Data Fig. 4g). Finally, we constructed a network that linked ligands, TFs, and ARGs together (Methods). Notably, the subnetwork of CEBPA is particularly interesting (Fig. 4g): CEBPA was proposed as a COVID-19 therapeutic target^46^ and identified as a key regulator in CD14 monocytes of hospitalized COVID-19 patients from scATAC-seq data^6^. Furthermore, 29/30 of its upstream ligands were either therapeutic targets or altered during SARS-CoV-2 infection, and 20/27 of its downstream ARGs were associated to COVID-19 or viral infection^46–53^. Two ARGs, SOCS3/SOCS3-DT and CAPN1, were potential targets for COVID-19 treatment^54^. Using MOCHA’s differential accessibility and co-accessibility modules, we constructed a putative upstream regulatory network that could be driving alternative TSS regulation in CD16 monocytes during early SARS-CoV-2 infection. Given that these results are largely aligned with the literature, we anticipate that this approach can be used more generally to identify potentially novel biological mechanisms.

### Longitudinal analysis of chromatin accessibility during COVID-19 recovery

To understand chromatin regulation during COVID-19 recovery, we analyzed scATAC-seq data of CD16 monocytes from our longitudinal COVID-19 study (Fig. 5a). The dataset, denoted as COVID19L, was collected on 69 longitudinal PBMC samples from 18 COVID+ participants (10 females and 8 males) over a period of 1–121 days PSO (Methods). We integrated MOCHA with existing tools and developed customized approaches to analyze the data at both single-cell and pseudo-bulk levels.

**Figure 5.**
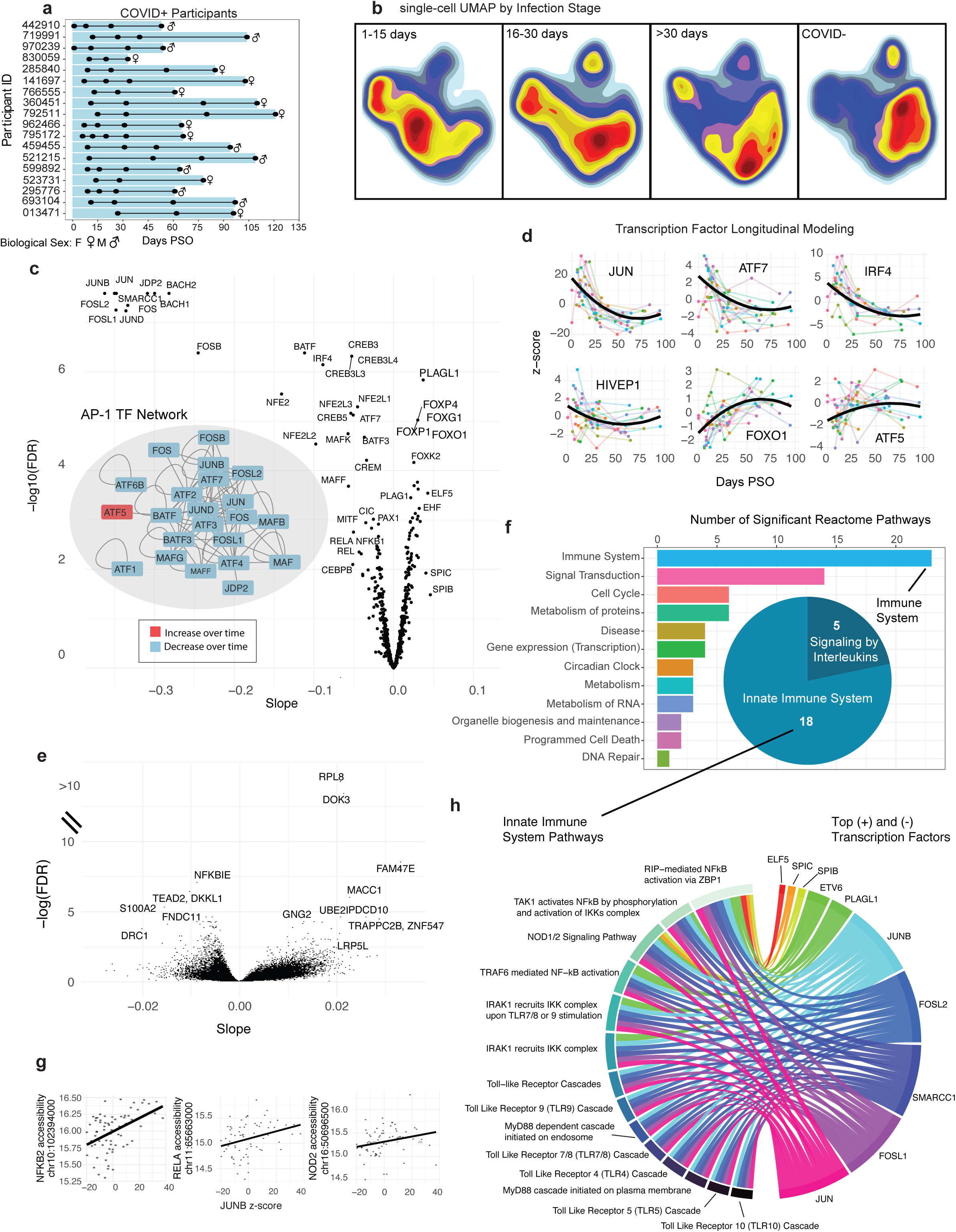
Integrative analyses to reveal longitudinal dynamics in CD16 monocytes during COVID-19 recovery. **a,** Longitudinal COVID19 cohort overview (n=18). Time points indicated by black dots illustrate sample availability for each COVID+ participant. **b,** Single-cell UMAP generated with tiles from the TSAM of CD16 monocytes in the COVID19 dataset (n=91 samples). Density plots were generated on COVID+ samples during early infection (1–15 days PSO, n=21), late infection (16–30 days PSO, n=13), and recovery (>30 days PSO, n=35), and on COVID- samples (n=22). **c,** Volcano plot illustrating the -log10(FDR) vs. the slope of motif usage over time. Motif usage was generated by running ChromVAR on tiles in the TSAM and extracting the corresponding z-scores. Insert: The network showing interacting TFs within the AP-1 family (APID database^87^) in which TFs were color-coded by the sign of their slope. **d,** Longitudinal motif usage over time for an exemplary set of TFs. Data of individual participants are shown in thin colored lines. The population trend is shown with a thick black line. **e,** Volcano plot illustrating the -log10(FDR) vs. the slope of gene promoter accessibility over time based on ZI-GLMM. A subset of the top promoters are labeled with their corresponding genes. **f,** Significant Reactome pathways (FDR < 0.1) enriched with genes having significant promoter accessibility changes. The pathways were aggregated into upper-level pathway annotations using Reactome’s database hierarchy. The barplot shows the number of pathways in each category. The pie chart breaks down the immune system pathways by Reactome’s next level categories. **g,** Three scatter plots illustrating examples of associations between gene promoter accessibility (y-axis) and JUNB’s ChromVar z-score (x-axis). The thick black line shows the population trend from ZI-GLMM. **h,** Bipartite network illustrating associations between the top 5 motifs having the largest positive (+) or negative (-) slopes and 14 significant innate immune pathways. For a link to be shown, motif-promoter associations must be found in at least 33% of significantly changing genes in the corresponding pathway. **c-h,** Based on data on CD16 monocytes in the COVID19L dataset (n=69). PSO: post symptom onset; UMAP: Uniform Manifold Approximation and Projection; TSAM: tile-sample accessibility matrix; FDR: false discovery rate; TF: transcription factor; AP-1: activator protein-1; ZI-GLMM: zero-inflated generalized linear mixed model. Source data are provided in Source Data Fig. 5. Figures were generated using Adobe Illustrator.

First, open tiles from the TSAM of CD16 monocytes were imported into ArchR as a peakset for dimensionality reduction. The resulting Uniform Manifold Approximation and Projection^55^ (UMAP) plot showed a clear shift in cellular population from initial infection to 30+ days post infection, at which time the cellular population appeared similar but not identical to that of the 22 COVID- participants (Fig. 5b).

Second, days PSO were binned (Fig. 5b) and used to learn a Monocle3^56^ trajectory, which largely followed the cellular population shift across the UMAP space (Supplementary Fig. 7a). Using ArchR, we further identified genes with GeneScore changes or tiles from the TSAM with accessibility shifts along this trajectory (Supplementary Fig. 7b). The genes with promoter accessibility shifts in CD16 monocytes were enriched in 72 immune system pathways, including 39 innate immune, 15 adaptive immune, and 18 cytokine signaling pathways (Supplementary Fig. 7c, right panel). In comparison, the corresponding pathway counts from the GeneScore analysis was only 10, 2, 4, and 4, respectively (Supplementary Fig. 7c, left panel). TSAM-based results were more informative and aligned better with the expected roles of CD16 monocytes than GeneScore-based ones.

Third, we converted the TSAM of CD16 monocytes into a chromVAR object and calculated sample-specific z-scores for TF activity (Methods). This enabled us to apply generalized linear mixed models (GLMM) to identify TFs with dynamic activities in CD16 monocytes during COVID-19 recovery. Among the 223 TFs whose activity changed significantly in time (FDR < 0.1, Fig. 5c-d), the AP-1 family (such as ATF1-7, JUN/B/D, MAF/F/G/K, FOS/B, and BACH1-2; Fig. 5c, insert) and the NF-κB family (such as REL/A and NFKB1-2) mostly had decreased activities, in consistency with their inflammatory, infection-responsive functions. On the contrary, the forkhead box (FOX) TF family (such as FOXP1/4, FOXG1, FOXO1, and FOXK2) had increased activities, which agrees with their known roles in immune homeostasis^57–59^. In comparison, we also identified 86 TFs with chromVAR z-score changes along the pseudotime trajectory described above, among which only 31 were unique (Supplementary Fig. 8). Longitudinal analysis based on real time identified more TFs with dynamic activities during COVID-19 recovery than the trajectory analysis based on pseudotime.

Fourth, the TSAM of CD16 monocytes was used to examine how gene promoter accessibility shifted during COVID-19 recovery. Since the data had about 20% zeros, we applied ZI-GLMM to address this specific challenge (Methods). A total of 2,120 genes demonstrated promoter accessibility shifts over time (FDR < 0.1), including genes regulating immune inflammation such as NFKBIE and DOK3 (Fig. 5e, Supplementary Fig. 9a)^60–62^. This gene set was enriched for 71 Reactome pathways (FDR < 0.1; Fig. 5f). Interestingly, among the 23 immune system pathways, five involve signaling by interleukins and 18 are related to the innate immune system (such as TLR-, MyD88-, and IRAK1-related pathways), but none are specific to the adaptive immune system. Again, these results are consistent with the expected roles of CD16 monocytes during viral infection.

Finally, to discover possible TF-gene regulations in CD16 monocytes during COVID-19 recovery, we applied ZI-GLMM to examine whether TFs with significant activity changes were associated with genes with significant promoter accessibility changes (Methods). For example, we found that JUNB chromVAR z-score was significantly associated (*P* < 0.05) with the promoter accessibility of 19 genes in the TLR4 cascade pathway (Fig. 5g, Supplementary Fig. 9b). As an illustration, we selected the top five TFs having the largest positive (PLAGL1, ELF5, ETV6, SPIB, and SPIC) or negative (JUN, FOSL1, SMARRC1, FOSL2 and JUNB) slopes, respectively, against time and examined their associations with gene promoters within the 18 innate immune pathways enriched during COVID-19 recovery (Fig. 5f). The five TFs having negative slopes were significantly associated (*P* < 0.05) with 14 pathways (Fig. 5h), consistent with the prominent roles of AP-1 TF family in innate immunity. Among the five TFs with positive slopes, PLAGL1 and ETV6 were significantly associated (p < 0.05) with five innate immune pathways, SPIB and SPIC with 2, and ELF1 with 1, respectively. Four innate immune pathways did not show significant associations with any of these top 10 TFs. While wet-lab validation is needed, such TF-gene associations generate interesting biological hypotheses regarding gene expression regulation in CD16 monocytes during COVID-19 recovery. Together, these results demonstrate MOCHA is a valuable tool in studying chromatin dynamics and gene regulatory networks based on longitudinal scATAC-seq data.

## Discussion

We developed MOCHA to robustly infer active gene regulatory programs in human disease cohorts based on scATAC-seq. First, we showed that our open chromatin model significantly agreed and was more sensitive in detecting sample-specific open chromatin regions than MACS2 and HOMER. Second, we identified differential accessible regions that better distinguish between COVID+ and COVID- participants than those by ArchR and/or Signac and uniquely revealed pathways affected by SARS-CoV-2 infection. Third, we constructed ligand-TF-gene networks on potential alternative TSS regulations during SARS-CoV-2 infection using only scATAC-seq data. Fourth, using zero-inflated mixed models, we identified motifs and promoters that were associated with COVID-19 recovery and constructed a TF-pathway network to infer which pathways were functionally important during COVID-19 recovery. MOCHA substantially increased the value of scATAC-seq in our COVID19 cohort by enabling robust modeling and visibility into the functional implications of chromatin accessibility. In addition, we illustrated how MOCHA can be integrated with existing tools such as ArchR, Monocle3, chromVAR, and NicheNet while enabling customized analysis using ZI-mixed effects models to gain unique insights from scATAC-seq data. Given its capabilities, we believe MOCHA is a valuable addition for analyzing scATAC-seq data, especially in biomedical research.

Constructing robust regulatory networks begins with reliable identification of patient- and cell type-specific open chromatin. We used peaks called by MACS2 on pseudo-bulk ATAC-seq data as imperfect “ground truth” to train and validate MOCHA. The training data of NK cells (n=179,836, 750 million fragments) had enough sequencing depth for reliable MACS2 performance and thus likely reliable MOCHA training. However, MACS2 might call every fragment as a peak for less abundant cell types, leading to many false positives. To mitigate such artifact, some pipelines artificially limit the number of peaks called by MACS2^11^. To provide a reasonable comparison, we focused our benchmarking on cell types with moderate to high cell counts. MOCHA outperformed MACS2 in calling sample-specific regions despite relying on MACS2 for training. In theory, MOCHA was not designed to call open tiles on datasets of mixed cells from multiple studies. For example, we used a global prefactor *S* to account for differences in data quality instead of, more properly, estimating an *S* for each of the many studies within the Hematopoiesis dataset. Nevertheless, MOCHA outperformed MACS2 and HOMER on all three datasets of varying data quality, although only slightly on the Hematopoiesis dataset. It is possible that the LRMs in MOCHA may need to be retrained if the difference in species, sample type, experimental protocol, sequencing depth, data quality, etc., becomes overwhelmingly large between our training data and user data. Due to a lack of access to GPU hardware^63^ and integration challenges^13^, we benchmarked MOCHA only with MACS2 and HOMER, which are the most widely incorporated open source peak callers. Given its strong performance against these standard algorithms, MOCHA’s robust open chromatin results provided a solid foundation for downstream analysis.

Additionally, gene regulatory networks require clear identification of accessibility changes. However, the presence of drop-out leads to many unreliable results. The incorporation of ZI statistical methods to handle drop out is a major advantage of MOCHA over existing tools. ZI methods provide well-documented improvements over their counterparts on ZI data^64,65^. While ZI methods are applied to scRNA-seq data exclusively at single-cell level, the sparsity of scATAC-seq data makes it necessary to apply ZI methods even at pseudo-bulk level. We applied the two-part Wilcoxon model^31^ for DAA, ZI correlation^33^ for CAA, and ZI-GLMM^30^ for longitudinal modeling in MOCHA and demonstrated how MOCHA led to more informative results than existing tools.

While single cell analysis provides granularity into cellular behavior, human cohort studies are usually interested in identifying patient-level behavior across cell populations. While current methods are centered at the single cell level, MOCHA aggregates scATAC-seq data into TSAMs to facilitate sample-centric analysis. To the best of our knowledge, this rather simple approach has not been reported to analyze scATAC-seq data. The approach provides several important advantages. First, the approach specifically addresses pseudo-replication bias in single-cell data and avoids computationally expensive single-cell mixed effect models, following recent advice for analyzing scRNA-seq data^28,29^. Second, the sample-centric approach makes it computationally feasible to analyze large, diverse human cohorts and explicitly models patient-level heterogeneity. Third, since the TSAM is constructed from standard Bioconductor data structures, its flexibility enables a broad range of scientific enquiries into gene and chromatin regulation and supports seamless integration with a variety of bioinformatics tools. For example, we applied ZI-GLMM and chromVAR to study COVID-19 recovery on our longitudinal scATAC-seq data. We believe TSAMs facilitate the extraction of genomic insights from large-scale, heterogeneous scATAC-seq data. Nevertheless, the approach is underpowered for studies of small sample size and not appropriate for comparing a handful of samples. We plan to adapt MOCHA for small-scale studies in the future.

We selected CD16 monocytes in our COVID19 dataset to showcase the utility of MOCHA in biomedical research. Our results reveal from multiple perspectives that the genomic regions associated with innate immune pathways (such as TLR, MyD88, and NF-κB) played essential roles in SARS-CoV-2 infection and patient recovery, aligning with the expected functions of CD16 monocytes during viral infection^66^. To the best of our knowledge, explicit longitudinal analysis on scATAC-seq data has not been reported, limiting the value of scATAC-seq in studying the regulatory landscapes of disease progression and recovery. Furthermore, despite the large number of publications on COVID-19, alternative TSS regulation during SARS-CoV-2 infection has not been reported. We consider MOCHA as a tool to generate interesting hypotheses from scATAC-seq data, which nevertheless need to be validated in follow-up studies. An in-depth, comprehensive analysis of our COVID19 cohort is beyond the scope of current work and will be presented in a follow-up paper.

In short, we present MOCHA as a tool to better infer gene regulation from scATAC-seq in biomedical and biological research. MOCHA is freely available as an R package in CRAN (https://cran.r-project.org/web/packages/MOCHA/index.html).

## Methods

### Longitudinal COVID-19 cohort

We recruited in the greater Seattle area n=18 participants (10 females and 8 males, aged 22–79 years) who tested positive (COVID+) for SARS-CoV-2 virus (Wuhan strain) and n=23 uninfected (COVID-) participants (10 females and 13 males, aged 29–77 years) for our longitudinal COVID-19 study^35^, “Seattle COVID-19 Cohort Study to Evaluate Immune Responses in Persons at Risk and with SARS-CoV-2 Infection”. All COVID+ participants had mild to moderate symptoms. Peripheral blood mononuclear cell (PBMC) and serum samples were collected from the COVID- participants at a single time point and from the COVID+ participants at 3-5 time points over a period of 1–121 days post-symptom-onset (PSO, total samples n=70). Study data were collected and managed using REDCap electronic data capture tools hosted at Fred Hutchinson Cancer Research Center (FHCRC). The FHCRC Institutional Review Board (IRB) approved the studies and procedures. Informed consent was obtained from all participants at the Seattle Vaccine Trials Unit to participate in the study and to publish their corresponding research data. Two participants declined to publish their raw sequencing data.

### COVID19 Single-cell ATAC-seq

#### PBMC isolation

Blood collected in acid citrate dextrose tubes was transferred to Leucosep tubes (Greiner Bio One). The tube was centrifuged at 800–1000 x g for 15 minutes and the PBMC layer recovered above the frit. PBMCs were washed twice with Hanks Balanced Solution without Ca+ or Mg+ (Gibco) at 200–400 x g for 10 min, counted, and aliquoted in heat-inactivated fetal bovine serum with 10% dimethylsulfoxide (DMSO, Sigma) for cryopreservation. PBMCs were cryopreserved at -80°C in Stratacooler (Nalgene) and transferred to liquid nitrogen for long-term storage.

#### FACS neutrophil depletion

To remove dead cells, debris, and neutrophils prior to scATAC-seq, PBMC samples were sorted by fluorescence-activated cell sorting (FACS) prior to cell permeabilization as described previously^67^. Cells were incubated with Fixable Viability Stain 510 (BD, 564406) for 15 minutes at room temperature and washed with AIM V medium (Gibco, 12055091) plus 25 mM HEPES before incubating with TruStain FcX (BioLegend, 422302) for 5 minutes on ice, followed by staining with mouse anti-human CD45 FITC (BioLegend, 304038) and mouse anti-human CD15 PE (BD, 562371) antibodies for 20 minutes on ice. Cells were washed with AIM V medium plus 25 mM HEPES and sorted on a BD FACSAria Fusion. A standard viable CD45+ cell gating scheme was employed: FSC-A x SSC-A (to exclude sub-cellular debris), two FSC-A doublet exclusion gates (FSC-W followed by FSC-H), dead cell exclusion gate (BV510 LIVE/DEAD negative), followed by CD45+ inclusion gate. Neutrophils (defined as SSChigh, CD15+) were then excluded in the final sort gate. An aliquot of each post-sort population was used to collect 50,000 events to assess post-sort purity.

#### Sample preparation

Permeabilized-cell scATAC-seq was performed as described previously^67^. A 5% w/v digitonin stock was prepared by diluting powdered digitonin (MP Biomedicals, 0215948082) in DMSO (Fisher Scientific, D12345), which was stored in 20 μL aliquots at −20°C until use. To permeabilize, 1×106 cells were added to a 1.5 mL low binding tube (Eppendorf, 022431021) and centrifuged (400 × g for 5 minutes at 4°C) using a swinging bucket rotor (Beckman Coulter Avanti J-15RIVD with JS4.750 swinging bucket, B99516). Cells were resuspended in 100 μL cold isotonic Permeabilization Buffer (20 mM Tris-HCl pH 7.4, 150 mM NaCl, 3 mM MgCl2, 0.01% digitonin) by pipette-mixing 10 times, then incubated on ice for 5 minutes, after which they were diluted with 1 mL of isotonic Wash Buffer (20 mM Tris-HCl pH 7.4, 150 mM NaCl, 3 mM MgCl2) by pipette-mixing five times. Cells were centrifuged (400 × g for 5 minutes at 4°C) using a swinging bucket rotor, and the supernatant was slowly removed using a vacuum aspirator pipette. Cells were resuspended in chilled TD1 buffer (Illumina, 15027866) by pipette-mixing to a target concentration of 2,300-10,000 cells per μL. Cells were filtered through 35 μm Falcon Cell Strainers (Corning, 352235) before counting on a Cellometer Spectrum Cell Counter (Nexcelom) using ViaStain acridine orange/propidium iodide solution (Nexcelom, C52-0106-5).

#### Tagmentation and fragment capture

scATAC-seq libraries were prepared according to the Chromium Single Cell ATAC v1.1 Reagent Kits User Guide (CG000209 Rev B) with several modifications. 15,000 cells were loaded into each tagmentation reaction. Permeabilized cells were brought to a volume of 9 μl in TD1 buffer (Illumina, 15027866) and mixed with 6 μl of Illumina TDE1 Tn5 transposase (Illumina, 15027916). Transposition was performed by incubating the prepared reactions on a C1000 Touch thermal cycler with 96– Deep Well Reaction Module (Bio-Rad, 1851197) at 37°C for 60 minutes, followed by a brief hold at 4°C. A Chromium NextGEM Chip H (10x Genomics, 2000180) was placed in a Chromium Next GEM Secondary Holder (10x Genomics, 3000332) and 50% Glycerol (Teknova, G1798) was dispensed into all unused wells. A master mix composed of Barcoding Reagent B (10x Genomics, 2000194), Reducing Agent B (10x Genomics, 2000087), and Barcoding Enzyme (10x Genomics, 2000125) was then added to each sample well, pipette-mixed, and loaded into row 1 of the chip. Chromium Single Cell ATAC Gel Beads v1.1 (10x Genomics, 2000210) were vortexed for 30 seconds and loaded into row 2 of the chip, along with Partitioning Oil (10x Genomics, 2000190) in row 3. A 10x Gasket (10x Genomics, 370017) was placed over the chip and attached to the Secondary Holder. The chip was loaded into a Chromium Single Cell Controller instrument (10x Genomics, 120270) for GEM generation. At the completion of the run, GEMs were collected and linear amplification was performed on a C1000 Touch thermal cycler with 96–Deep Well Reaction Module: 72°C for 5 min, 98°C for 30 sec, 12 cycles of: 98°C for 10 sec, 59°C for 30 sec and 72°C for 1 min.

#### Sequencing library preparation

GEMs were separated into a biphasic mixture through addition of Recovery Agent (10x Genomics, 220016); the aqueous phase was retained and removed of barcoding reagents using Dynabead MyOne SILANE (10x Genomics, 2000048) and SPRIselect reagent (Beckman Coulter, B23318) bead clean-ups. Sequencing libraries were constructed by amplifying the barcoded ATAC fragments in a sample indexing PCR consisting of SI-PCR Primer B (10x Genomics, 2000128), Amp Mix (10x Genomics, 2000047) and Chromium i7 Sample Index Plate N, Set A (10x Genomics, 3000262) as described in the 10x scATAC User Guide. Amplification was performed in a C1000 Touch thermal cycler with 96–Deep Well Reaction Module: 98°C for 45 sec, for 9 to 11 cycles of: 98°C for 20 sec, 67°C for 30 sec, 72°C for 20 sec, with a final extension of 72°C for 1 min. Final libraries were prepared using a dual-sided SPRIselect size-selection cleanup. SPRIselect beads were mixed with completed PCR reactions at a ratio of 0.4x bead:sample and incubated at room temperature to bind large DNA fragments. Reactions were incubated on a magnet, and the supernatant was then transferred and mixed with additional SPRIselect reagent to a final ratio of 1.2x bead:sample (ratio includes first SPRI addition) and incubated at room temperature to bind ATAC fragments. Reactions were incubated on a magnet, the supernatant containing unbound PCR primers and reagents was discarded, and DNA bound SPRI beads were washed twice with 80% v/v ethanol. SPRI beads were resuspended in Buffer EB (Qiagen, 1014609), incubated on a magnet, and the supernatant was transferred resulting in final, sequencing-ready libraries.

#### Quantification and sequencing

Final libraries were quantified using a Quant-iT PicoGreen dsDNA Assay Kit (Thermo Fisher Scientific, P7589) on a SpectraMax iD3 (Molecular Devices). Library quality and average fragment size were assessed using a Bioanalyzer (Agilent, G2939A) High Sensitivity DNA chip (Agilent, 5067-4626). Libraries were sequenced on the Illumina NovaSeq platform with the following read lengths: 51nt read 1, 8nt i7 index, 16nt i5 index, 51nt read 2.

#### Data preprocessing

scATAC-seq libraries were processed as described previously^67^. In brief, cellranger-atac mkfastq (10x Genomics v1.1.0) was used to demultiplex BCL files to FASTQ. FASTQ files were aligned to the human genome (10x Genomics refdata-cellranger-atac-GRCh38-1.1.0) using cellranger-atac count (10x Genomics v1.1.0) with default settings. Fragment positions were used to quantify reads overlapping a reference peak set (GSE123577_pbmc_peaks.bed.gz from GEO accession GSE123577^68^), which was converted from hg19 to hg38 using the liftOver package for R^69^, ENCODE reference accessible regions (ENCODE file ID ENCFF503GCK^70^), and TSS regions (TSS ±2kb from Ensembl v93^71^ for each cell barcode using a bedtools (v2.29.1 ^72^) analytical pipeline.

#### Quality control

Custom R scripts were used to remove cells with less than 1,000 uniquely aligned fragments, less than 20% of fragments overlapping reference peak regions, less than 20% of fragments overlapping ENCODE TSS regions, and less than 50% of peaks overlapping ENCODE reference regions. The ArchR package^11^ was used to assess doublets in scATAC data. Doublets were identified using the ScoreDoublets function using a filter ratio of 8, and cells with a Doublet Enrichment score exceeding 1.3 as determined by ArchR’s doublet detection algorithm^11^ were not considered for downstream analysis.

#### Dimensionality reduction and cell type labelin

We used the ArchR package to generate a count matrix for a PBMC reference peak set^68^. Dimensionality reduction was performed using the ArchR addIterativeLSI function (parameters varFeatures = 10,000, iterations = 2), and the addClusters function was used to identify clusters in latent semantic indexing (LSI) dimensions using the Louvain community detection algorithm. For visualization, Uniform Manifold Approximation and Projection^55^ (UMAP) was performed using ArchR’s addUMAP function at the default settings. The ArchR addGeneIntegrationMatrix function (parameters transferParams = list(dims = 1:10, k.weight = 20)) was used to label scATAC cells using the Seurat level 1 cell types from the Seurat v4.0 PBMC reference dataset^73^. To generate clusters that more closely matched label transfer results, we performed K-means clustering on the UMAP coordinates using 3 to 50 cluster centers and identified a set of clusters that each had > 80% of cells sharing a single cell type identity. Almost all such clusters contained >= 98% cells from a single major cell type (T cells, B cells, NK cells, or monocytes/DCs/other), with the exception of a single cluster with 88% purity. We used clusters of the same major cell type to subset the data into T cells, B cells, NK cells, or monocytes/DCs/other for downstream analyses. For each major cell type, we repeated the same dimensionality reduction (LSI/UMAP) process on the scATAC-seq data with the same settings. We then performed a second round of label transfer, using the ArchR addGeneIntegrationMatrix function (same parameters as described above for level 1), to reach level 2 and 3 cell labelings of the Seurat PBMC reference dataset. These labels were consolidated into 25 cell types for most analysis, except for the co-accessibility analysis where 17 cell types were used to match the published promoter-capture HiC resource^43^. The median cell labeling score across all cells that passed quality control was 0.74.

### Three scATAC-seq datasets for MOCHA development and benchmarking

#### COVID19 dataset

Two samples (1 COVID- sample, male; 1 COVID+ sample, female, collected on day 12 PSO) from our longitudinal COVID-19 cohort were lost due to low sample volume. The scATAC-seq data of the remaining samples was denoted as the COVID19 dataset (n=91) in this study. After removing doublets and cells of poor quality, high quality data of 1,311,638 cells were obtained. The data was split into two overlapping subsets for some analyses: 1) A cross-sectional dataset (denoted as COVID19X, n=39) included data of COVID- samples (n=22, 10 females and 12 males) and the first samples of COVID+ participants (n=17, 9 females and 8 males) during early infection (<16 days PSO). 2) A longitudinal dataset (denoted as COVID19L, n=69) included all data for the 18 COVID+ participants (10 females and 8 males). The overlap between COVID19X and COVID19L was 17, which were the first samples of COVID+ participants. The full dataset (COVID19) can be accessed at GEO under accession number GSE173590. (Note to reviewers: data will be released to the public prior to the first publication of our manuscripts.)

#### HealthyDonor dataset

This longitudinal scATAC-seq dataset^39^ was collected on 18 PBMC samples of 4 healthy donors (aged 29–39 years) over 6 weeks (1 female and 1 male, weeks 2–7; 2 males, weeks 2, 4, and 7). The donors had no diagnosis of active or chronic disease during the study. The data is publicly available at GEO under accession number GSE190992. We used the dataset as is, except we removed cells with doublet enrichment score exceeding 1.3, based on ArchR’s doublet detection algorithm^11^. High quality data of 145,711 cells were obtained. From this dataset, we consolidated existing annotations into 25 cell types with a published median cell labeling score of 0.78.

#### Hematopoiesis dataset

This dataset was downloaded from (https://www.dropbox.com/s/sijf2votfej629t/Save-Large-Heme-ArchRProject.tar.gz). It consists of ∼220,000 hematopoietic cells from the hematopoiesis dataset in ArchR^11^. As described in their Supplementary Table 1, the dataset was assembled from 49 samples in four data sources, of different sample types (mixed, sorted, and unsorted cells; PBMCs; and bone marrow mononuclear cells), and generated using different sample processing protocols and on different technical platforms. We used ArchR to generate doublet scores and removed clusters with both high doublet scores and a mixture of disparate cell types. This doublet removal only applied to sequencing wells that were not sorted, purified cells. In the end, data of 95,599 cells were obtained. Because many of the cell types were sorted populations run on an individual well, many cell types were not available across samples. As a result, we treated all cell types as coming from a single sample for benchmarking purposes. We used previously published cell annotations, with a median labeling score of 0.70.

### Assessing Dataset Noise using the Altius Peakset

To assess ‘noise’ within a dataset (i.e., fragments from closed regions known as heterochromatin), we used the Altius consensus peakset ^70^ of over 3.6 million DNase I hypersensitive sites within the human genome as an approximation of all potentially accessible sites. We calculated the overlap rate between fragments and the Altius consensus peakset for each cell type and dataset, in order to assess the quality of the data. The COVID19 dataset had an median Altius peakset overlap rate of 88.9%, while the corresponding rates for the Healthy Control dataset and the Hematopoiesis dataset were 75.5% and 82%, respectively.

### MOCHA overview

MOCHA is implemented as an open-source R package under the GPLv3 license in CRAN (https://CRAN.R-project.org/package=MOCHA). All code and development versions of MOCHA are available at https://github.com/aifimmunology/MOCHA.

MOCHA is designed to run in-memory and interoperate with common Bioconductor methods and classes (e.g., RaggedExperiment, MultiAssayExperiment, and Summarized Experiment). It takes as input four objects that are commonly generated from scATAC-seq after cell labeling and the removal of doublets and cells of low quality data. These four objects are: 1) a list of GRanges or GRangesList containing per-sample ATAC fragments, 2) cell metadata with cell labels, 3) a BSGenome annotation object for the organism, and 4) a GRanges containing blacklisted regions. These inputs can be passed to MOCHA directly from an ArchR object. Alternatively, results can be extracted from Signac, SnapATAC, or ArchR, and converted to common Bioconductor data objects, which can then be imported into MOCHA. By operating on well-supported Bioconductor objects, MOCHA’s inputs and outputs are compatible with the broader R ecosystem for sequencing analyses, and are easily exportable to genomic file formats such as BED and BAM.

MOCHA’s core functionality runs as a pipeline from these inputs to perform sample-specific open tile prediction and consensus analysis, resulting in a TSAM represented as a Bioconductor RangedSummarizedExperiment. On systems with sufficient memory, MOCHA’s functions can be parallelized over samples with the ‘numCores’ parameter to decrease runtime. From the TSAM, MOCHA provides functions for zero-inflated (ZI) co-accessibility and ZI differential accessibility analysis. The format of the TSAM output enables additional downstream analyses with other R packages. For example, the TSAM RangedSummarizedExperiment can be used directly as the counts matrix input for motif deviations analysis with chromVAR (https://greenleaflab.github.io/chromVAR/articles/Articles/Counts.html). Additional details on the workflow and functions on the MOCHA package are provided in Supplemental Fig. 1.

### Tiling the Genome

MOCHA splits the genome into pre-defined, non-overlapping 500 base-pair tiles that remain invariant across samples and cell types. MOCHA annotates each tile using a user-provided transcript database (e.g., HG38 Transcript Database) as follows: Promoter regions are 2000 bp upstream and 200 bp downstream from transcriptional start sites^74^. Intragenic regions are tiles that fall within a gene body, but not within the promoter regions. All other regions are classified as distal.

From there, we only consider tiles that overlap with ATAC fragments. MOCHA counts the number of fragments in a tile as follows

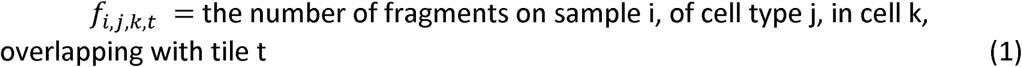

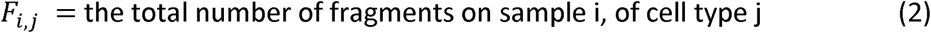

If a fragment falls between two tiles, it is counted on both tiles.

### Normalization

#### Normalization Techniques Using Invariant CTCF Sites

We examined three normalizing approaches: dividing the number of fragments by 1) the total number of fragments for sample i, cell type j (i.e.,*F_i,j_*); 2) the total number of fragments for sample i (i.e., *F_i,j_* = *Σ_j_F_i,j_*), and 3) the total number of cells in sample i, cell type j. We evaluated the above normalization methods along with the raw data based on a list of 2230 cell-type invariant CCCTC-binding factor (CTCF) sites from the ChIP Atlas database^75^. These loci were identified in at least 201/204 (99%) of blood cell types present in the ChIP-seq Atlas database ^76^. Using these CTCF sites, each approach was assessed based on the corresponding distribution of coefficient of variation (CV) in peak accessibility. MOCHA normalizes data using *F_i,j_*

#### Sample- and cell type-specific Normalization

For each sample i, cell type j, and tile t, MOCHA calculates the following normalized features:

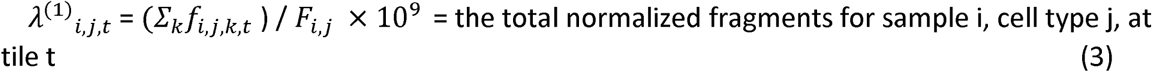

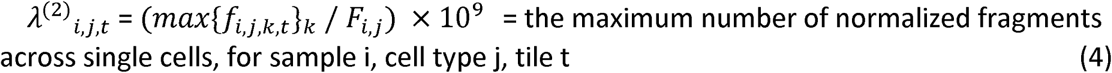

Since the NK population used for model training contained 750 million fragments, a scaling factor of 10^9^ is applied to make the raw and normalized counts on the same scale across cellular abundances, and keep normalized values greater than 1 to minimize convergence errors in downstream model training. Biologically, 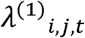 is designed to capture the total number of fragments across all cells (e.g., pseudo-bulk), normalized by the sequencing depth for that cell type and sample. Given the sparsity of scATAC-seq data and the assumption of limited number of genomic copies (2x-4x) in a typical cell, 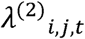 is designed to capture the presence of multiple fragments in a tile from any cell, which can only be evaluated on single cell data. This approach combines single cell and pseudo-bulk information for downstream prediction. Normalizing by *F_i,j_* is used to normalize both sequencing depth and cell population variability. This approach provides both a sample- and cell-type specific normalization scheme.

### Evaluation of open chromatin accessibility

#### Training of logistic regression models (LRMs) for predicting tile accessibility

MOCHA assumes a typical ploidy per cell (two to four copies of the genome). Its pseudocode and further details are provided in Supplementary Fig. 2 in order to allow for modifications when the above assumption does not hold.

We used scATAC-seq data of 179k NK cells in the COVID19 (n=91) dataset as the training dataset. First, we normalized the scATAC-seq data and collapsed it into pseudobulk data. Second, we applied MACS2 ^37^ (’-g hs -f BED --nolambda --shift -75 --extsize 150 --broad’, ‘--model -n’) to identify accessible peaks in the pseudobulk data, using previously published parameters for identifying peaks in scATAC-seq with the modification to call broad rather than narrow peaks. The resulting peaks were then overlaid onto our pre-defined 500 bp tiles. We trim the broad peaks by 75 base pairs at each end to remove the tail ends of peaks that may extend onto tiles with no signal. MACS2 identified 1.15 million tiles as ‘accessible regions’. We labeled all other fragment-containing regions as inaccessible, and used these ‘accessible’ and ‘inaccessible’ regions for training. Third, we randomly selected NK cells at cell counts ranging from 170k to 5 at discrete intervals, generating 10 replicates for subsets < 50k cells, and 5 replicates for larger subsets. In each of the subsets, we calculated 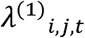 and 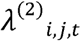 at individual tiles. Fourth, we trained a LRM for each selected subset of NK cells based on the tile labeling just described. For each sample of *n_a_* cells, the LRM calculates a probability score to assess the likelihood of a tile being accessible, using the formula

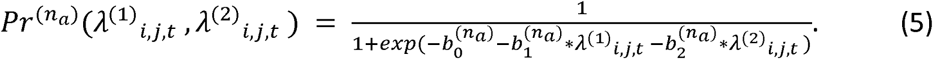

Here 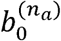 is the intercept, 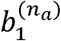 and 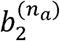 are coefficients for 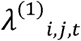 and 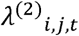 respectively. A tile is predicted as accessible if 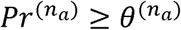 or inaccessible if 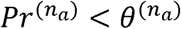 where 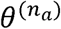 is the threshold value separating accessible and inaccessible tiles. We used Youden index^77^ to calculate 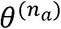 in this study, using the *cutpointR* R package.

Fifth, we collected 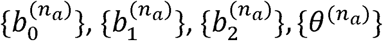 from the 10 or 5 replicated runs on *n* cells and then took the corresponding median coefficients, i.e., 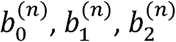, and *θ*^(*n*)^, to construct the predictive model for a sample of *n* cells. Finally, we used the learned coefficients and the learned thresholds to smoothen the model to interpolate the model across cellular abundances that the model was not trained on. The final model is composed of a set of smoothened coefficients 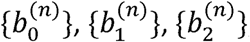, and smoothened thresholds{*θ*^(*n*)^} from all examined *n*.

#### Prediction of tile accessibility on new data

To predict accessibility in a new dataset, MOCHA first accounts for differences in sequencing depth and cell count across datasets and calculates the ratio, S, of the median (across samples) number of total fragments in the training data, and the corresponding median in the new dataset. MOCHA then scales both 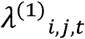 and 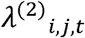 in the new dataset by S and calculates the likelihood of a tile being accessible as

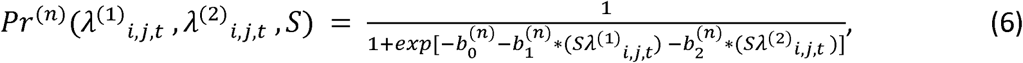

where *n* is the number of cells of the targeted cell type in the targeted sample. As before, a tile is predicted as accessible if 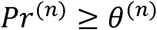 or inaccessible if 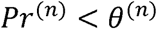 where *θ*^(*n*)^ is the threshold value separating accessible and inaccessible tiles.

#### Benchmarking Open Regions

To benchmark MACS2, HOMER, and MOCHA, we ran each tool per sample and cell type to generate comparable accessibility measurements across three cell types in three different datasets. For MACS2, we used the following parameters to call broad peaks (’-g hs -f BED --nolambda --shift -75 --extsize 150 --broad’--nomodel -n’), in accordance with previous published scATAC-seq settings^11^. For HOMER, we used the *findPeaks* function with default parameters, and added (’-style histone’) to call broad peaks. While HOMER and MACS2 are primarily designed around the properties of ChIP-seq and DNase-seq, they are also recommended for use with bulk ATAC-seq^78^.

To ensure head-to-head comparisons, we overlaid HOMER and MACS2’s peaks into MOCHA’s predefined 500 base-pair tiles to translate peak calls into open tile calls. Similar to training, we trimmed 75 bp off each end, as MACS2’s shift/extsize parameters extends fragments to improve peak calling under the -nomodel flag^79^. By trimming, we avoid counting the tails of peaks that might extend into tiles with no actual fragments. This trimming approach allows for a direct head-to-head comparison of open regions detected across methods. Additionally, all three methods are provided the same normalized pseudo-bulk intensity information to ensure comparable peak calling and prevent confounding peak calling and normalization. After translating MACS2 and HOMER peaks into open tiles, we then compared the number of open tiles per sample across all methods, cell types, and datasets.

Next, we generated a TSAM for each cell type across all three methods. The TSAM is a matrix with an array-type structure, where each cell contains the normalized *λ*^(1)^ intensities for a given sample i, at tile j. We kept open tiles that were called in at least 20% of samples (or all tiles in Hematopoiesis). By generating a TSAM for each method, we compared reproducible, population-level open tiles across all three methods. The 20% threshold was applied to filter out noisy data.

#### CTCF and TSS Sites for benchmarking

CTCF sites were drawn from the ChipSet Atlas^75^. In brief, we download a bed file containing CTCF peaks for all blood cell types, and then used Plyranges’s reduce_ranges^80^ function to collapse duplicate peak calls into one non-redundant and smaller file for detecting overlaps. This process was done for both Hg19 (n= 197,882) and Hg38 (n=184,588). TSS sites were taken from Bioconductor database TxDb.Hsapiens.UCSC.hg19.knownGene for the Hematopoiesis dataset (which was aligned to Hg19), and TxDb.Hsapiens.UCSC.hg38.refGene for the other datasets by first extracting the transcripts for all genes. The TSS were then extracted from the transcripts using the promoters() command (Hg19, n =62,265, Hg38, n= 88,819). We then calculated the number of tiles that overlapped with a CTCF and TSS site using the subsetByOverlaps function from the GenomicRanges^74^ R package.

#### Runtime Comparison on open chromatin analysis

Using the CD14 monocyte population in our COVID19 dataset (n=91), we produced 13 subsamples ranging from 100,000 to 10 cells and measured 10 replicates of the time it takes to conduct open chromatin analysis. Our runtime comparisons were conducted on N2 machines on the Google Cloud Platform, with 64 vCPUs and 512GB RAM. MOCHA version 0.2.0 was used. The R package “tictoc” was used to record elapsed time.

#### Downsampling Comparison on open chromatin

Since pooling cells across samples before calling open tiles is a common approach, we benchmarked all three methods on the same randomly selected cell subsets ranging from 5 cells to the full set in the COVID19 dataset (n=91). For this comparison, we utilized the same three cell types across the same three datasets. For each cell type and dataset, we used the following procedures. For MOCHA:

1. Generate a coverage object using predefined 500 base-pair tiles on all pooled cells (e.g., CD16 monocytes).
2. Predict open tiles on the pooled cells.
3. Count the total number of open tiles, the number of open tiles overlapping with CTCF sites, and the number of open tiles overlapping with TSSs.
4. Repeat (1–3) for all pre-specified downsampled cell counts.

For MACS2 and HOMER:

1. Generate a coverage file on all pooled cells (e.g., CD16 monocytes),
2. Call peaks on the pooled cells.
3. Convert the peak regions onto the pre-defined MOCHA tiles.
4. Count the total number of open tiles, the number of open tiles overlapping with CTCF sites, and the number of open tiles overlapping with TSSs.
5. Repeated (1–4) for the pre-specified downsampled cell counts.

### Differential Accessibility Analysis (DAA)

#### MOCHA’s zero-inflated method for DAA

MOCHA identifies differential accessibility tiles (DATs) in a targeted cell type between sample groups A and B in three steps:

First, similar to others ^10,11^, MOCHA prioritizes tiles for testing using heuristic functions to calculate two data-driven thresholds. MOCHA transforms the total fragment count 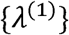 in the corresponding TSAM to 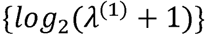 and fits a mixture model of two normal distributions on all 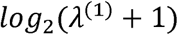 values in the TSAM (Supplementary Fig. 2g). This bimodal model provides a heuristic threshold to prioritize high-signal tiles. From there, we used the TSAM metadata to identify any differences in sequencing depth by comparing the median number of fragments per sample between groups. This analysis informs the ZI threshold. Given our initial observations of a 25% difference in fragment counts, we set a 50% threshold (2X the observed sequencing depth difference) to control for technical artifacts. Tiles that do not pass either threshold are assigned a DAT *P* value of NA, and those passing thresholds are then tested for differential accessibility.

Second, MOCHA tests for differential accessibility as follows. Denote the percentages of zeroes among samples of the two groups as *ρ_A_* and *ρ_B_* and the corresponding medians of non-zero 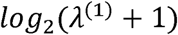 values as *μ_A_* and *μ_B_* MOCHA then tests whether a tile is a DAT based on the following hypothesis testing:

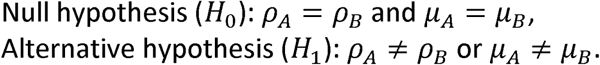

MOCHA uses the two-part Wilcoxon (TP-W) test^32^ to combine results from the binomial test on *ρ_A_* and *ρ_B_* with results from the Wilcoxon rank-sum test on *μ_A_* and *μ_B_*. Since each test statistic can be transformed to follow a 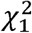 distribution (i.e., 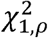 and 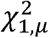), MOCHA combines them into a single test statistic, i.e., 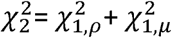, and consequently evaluates from it a single *P* value^31^. In the absence of zeros, the TP-W test mathematically reduces to the standard Wilcoxon rank-sum test.

Finally, to control for multiple testing, MOCHA evaluates a false discovery rate (FDR)^81^ for each tile and uses a default threshold of 0.2 to identify DATs. Since the *P* values are inflated near 1 (see Supplementary Fig. 5a), the background in the FDR calculation is estimated from *P* ≤ 0.95 only.

In addition, MOCHA uses the Hodges-Lehmann estimator^82^ to estimate Log2(fold change) on chromatin accessibility between the two sample groups. More specifically, MOCHA first calculates the difference between each sample pair (one sample each from group A or B) having non-zero 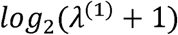 values and then takes the median from all paired differences as an estimate for Log2(fold change) between the two sample groups.

#### Benchmarking MOCHA with ArchR and Signac on DAA

ArchR and Signac’s DA modules were each run on a single cell count matrix generated from the same tile set (215,649 tiles) as the COVID19X CD16 monocytes TSAM. For ArchR, default settings were used, except we modified maxCells to include all cells (n = 24744). For Signac, we lowered the minimum percent detection (pct = 0.001), and the log2FC threshold (logfc.threshold = 0.05) in order to test the full tileset, thus enabling a full head-to-head comparison. As a close analog of Signac’s tutorial, we also set latent.vars to ‘nFrags’ to adjust for sequencing depth.

### Assessing Discriminative Power Per Method

We randomly subsampled 50 DATs from the output of each method, ran K-means clustering (K=2), and generated the following confusion matrix to summarize the predictions.

|  | COVID+ | COVID- |  |
| --- | --- | --- | --- |
| Cluster 1 | a | b | a+b |
| Cluster 2 | c | d | c+d |
|  | a+c | b+d | a+b+c+d |

We then calculated Holley’s^42^ 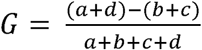 to assess how well the 50 randomly selected DATs in separating COVID+ and COVID- samples. We used |G| for the comparison since it is irrelevant which cluster is enriched for COVID+ samples. We repeated this process 1,000 times to obtain a distribution for each method.

#### DAA runtime comparison

To evaluate each method’s speed in DAA, we started by testing all 215,649 tiles in the CD16 monocytes TSAM, and gradually decreased the number of tested tiles. At each downsample, we tracked the run time required to identify DATs within those randomly selected tiles. The tictoc R package was used to calculate runtime.

### Co-Accessibility Analysis (CAA)

#### MOCHA’s zero-inflated (ZI) method for CAA

MOCHA applies ZI Spearman correlation^33,34^ to evaluate the co-accessibility of two tiles (e.g., *x* and *y*) across either cell types or samples based on the corresponding 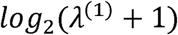 values. More specifically, the method first calculates the standard Spearman correlation on (*x*,*y*) pairs of non-zero data (i.e., *x* > 0 and *y* > 0), denoted as *ρ_S_*_,11_, and then adjusts it in the presence of zeroes in either tile as follows:

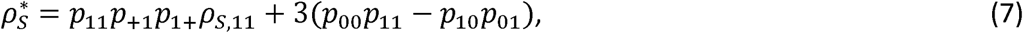

where

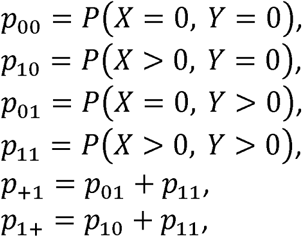

which quantify how zeros are distributed among the two tiles across all data points with *p*_00_ +*p*_10_ + *p*_01_ +*p*_11_. In the absence of zeros, the ZI-Spearman correlation reduces to the standard Spearman correlation, i.e., 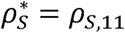. MOCHA makes two modifications to an R implementation of the method^25^: 1) The Spearman correlation (*ρ_S_*_,11_) is calculated in C language for optimal computing time and 2) undefined ZI Spearman correlations (when *ρ_S_*_,11_ cannot be calculated) are assigned to NA rather than replacing them with the standard Spearman correlations with zeros treated as normal data.

#### Benchmarking inter-cell-type co-accessibility

We c promoter-capture HiC (pcHiC) resource^43^ which identified promoter-enhancer regulatory links. From there, we used the liftOver R package, version 1.22.0, and the Hg19 to Hg38 conversion file (hg19ToHg38.over.chain, https://hgdownload.soe.ucsc.edu/gbdb/hg19/liftOver/) to convert promoter/enhancer loci from HG19 to HG38. Promoters and enhancers were then tiled into 500 bp windows to generate all promoter-enhancer tile (PET) pairs. We kept only PET pairs when both tiles were identified as accessible by MOCHA in naive CD4+ and CD8+ T Cells and had pcHiC evidence supporting their interaction specifically in naive CD4+ and CD8+ T cells. The obtained 1.2 million PET pairs were then treated as ‘ground truth’ for benchmarking the standard and the ZI Spearman methods in evaluating inter-cell-type co-accessibility. For comparison, we randomly selected 100k non-PET tile pairs across the genome as a negative background.

We applied both Spearman methods to calculate the inter-cell-type correlation values between both the PET and the random pairs, stacking data of different samples in the COVID19 dataset together (n=91). The inter-cell-type correlation was calculated across 17 cell types, including B intermediate, B memory, B naïve, CD14 Mono, CD16 Mono, CD4 Effector, CD4 Naïve, CD8 Effector, CD8 Naïve, DC, HSPC, MAIT, NK, NK Proliferating, NK_CD56bright, OtherT, and Treg. We used the Kolmogorov-Smirnov (KS) distance to quantify how well the two Spearman methods separated the PET pairs from the random pairs. To identify significant PET pairs by either Spearman method, we first treated the corresponding distribution of correlation values of the random pairs as a null distribution, calculated an empirical *P* value for each PET pair based on its correlation value, and then converted the obtained *P* values of all PET pairs into FDR values. A PET pair was considered as significant if FDR < 0.1.

### Pathway enrichment analysis

Pathway enrichment analysis was mostly restricted to the Reactome pathway database. All genes within the database reference of TxDb.Hsapiens.UCSC.hg38.refGene from Bioconductor^83^ were selected as the background. Over-representation analysis was performed using WebGestaltR^84^. We annotated enriched pathways at the highest level within the Reactome’s database hierarchy. Lower level annotations on immune system pathways were provided to discern adaptive, innate, and general signaling pathways. Using WebGestaltR, pathway enrichment analysis was performed once on Wikipathways, Gene Ontology (Biological Processes, Non-redundant), and KEGG for illustrative purposes.

### Identification of alternatively regulated transcription start sites (TSSs)

We extracted all TSSs from the Transcript database TxDb.Hsapiens.UCSC.hg38.refGene found on BioConductor ^83^, and then expanded them upstream by 125 bp to account for TSSs falling very close to a tile boundary. We filtered out genes with only one TSS. If alternative TSSs of the same gene occurred within a user-defined neighborhood (default: 150 bp) of each other, we collapsed them into a single TSS. We then found the intersection between alternative TSSs and the 6211 DATs between COVID+ and COVID- samples in CD16 monocytes. TSSs that landed on a DAT were assigned with the FDR of the corresponding DATs. We categorized alternatively regulated genes (ARGs) as

- Type I: A gene had a subset of TSSs showing differential accessibility (FDR < 0.2) in the same direction and another subset being open but not differential.
- Type II: A gene had at least two TSSs showing differential accessibility (FDR < 0.2) but in opposite directions.

### Motif enrichment analysis

Motif matching was done using the *motifmatchr* package and the CISBP motif database, as provided by the chromVARmotif package (https://github.com/GreenleafLab/chromVARmotifs). MOCHA uses a standard hypergeometric test to identify enriched motifs, with a user-provided foreground and background tile sets. For multi-testing corrections, the resulting p-values were converted into FDRs. To understand the upstream signaling mechanisms regulating ARGs, we first applied MOCHA to identify tiles that were within ±1M bp of and co-accessible (inter-sample, ZI-Spearman correlation > 0.5) with the corresponding DATs. These DATs and their co-accessible tiles were selected as the foreground tile set. For the background tile set, we chose all tiles with TSSs and their co-accessible tiles that did not overlap with the foreground set. These foreground and background tile sets were used to calculate CISBP motif enrichment regulating ARGs.

### Ligand-motif set enrichment analysis (LMSEA)

The NicheNet^45^ database has identified links between upstream ligands and downstream transcription factors (TFs) that regulate gene expression^45^. Using the same principle as pathway enrichment analysis, we designed a Ligand-Motif Set Enrichment Analysis (LMSEA) framework to capture potential drivers of our observed motifs (i.e., ligands regulate the TFs in our dataset). Specifically, LMSEA tests whether motifs linked to a ligand of interest are significantly (using hypergeometric test) over-represented in our observed motifs relative to the ligand’s motif set within NicheNet. The Benjamini and Hochberg (BH) procedure was used to adjust *P* values for multiple comparisons. An adjusted *P* value < 0.05 was considered significant.

### Construction of ligand-transcription factor-gene network

We constructed ligand-TF-gene networks and visualized them using Cytoscape^85^. The nodes were ARGs, enriched motifs (TFs), and enriched ligands. Edges were drawn as follows: a motif-gene link was created if an enriched TF was found within the TSS-containing DATs of an ARG or their co-accessible tiles, a ligand-motif link was drawn if a ligand was known to interact with a TF in NicheNet’s ligand-transcription matrix.

### Longitudinal analysis of COVID-19 response at single-cell level

#### Grouping COVID+ samples by infection stage

COVID+ samples (n=69) in the COVID19 dataset were grouped by the corresponding infection stage, including early infection (1–15 days PSO, n=21), late infection (16–30 days PSO, n=13), and recovery (>30 days PSO, n=35).

#### Generation of density UMAP

We extracted sample-specific open tiles on CD16 monocytes for all samples in the COVID19 dataset (n=91). From there, we generated a TSAM by aggregating all tiles that were called in at least 20% of samples at any infection stage or uninfected. We extracted the tiles from the resulting TSAM and added them to the original ArchR project via addPeakSet. We then generated a single-cell peak matrix from this tile set, using addPeakMatrix, and used it as input for ArchR’s iterative LSI and UMAP functions. The LSI was run with default parameters, except for the number of iterations (5 instead of 2). The UMAP was run on standard ArchR settings^11^ on the resulting iterative LSI object. Based on the resulting single-cell UMAPs, we generated a density plot for each infection stage or uninfected.

#### Pseudotime trajectory analysis

We used ArchR’s standard Monocle3 pipeline to conduct a trajectory analysis. We instructed Monocle to construct a trajectory from cells belonging to samples in the order of early infection, late infection, recovery, and uninfected. The resulting trajectory was overlaid on the single-cell UMAP. Following the above trajectory, three distinct pseudotime heatmaps were generated using ArchR’s standard protocol and the following input single-cell matrices: log2-normalized GeneScores, peak (tile) accessibility, and ChromVAR z-scores. Using ArchR’s functions with default settings, we further extracted pseudotime-changing elements for each of the three matrices.

### Longitudinal analysis of COVID-19 response at pseudo-bulk level

#### Longitudinal analysis of motif usage

We modeled longitudinal motif usage using pseudobulk ChromVar motif z-scores. We converted the TSAM of CD16 monocytes from the COVID19L dataset (n=69) into a ChromVAR-compatible object, and then ran ChromVAR on the TSAM-derived object to generate sample-level motif z-scores. We then modeled motif usage with the following generalized linear mixed effect model (GLMM):

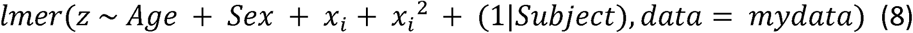

Where *x_i_* was the centered days PSO (i.e., days PSO of individual samples minus the mean days PSO of all samples)^86^ and (1|*Subject*) indicated that random intercepts were used for individual participants. The *P* values associated with the linear *x_i_* terms were extracted using the lmerTest package. We converted the *P* values to FDRs to control multiple testing. Motifs with a FDR < 0.1 were considered as significantly changing in time.

#### Transcription factor (TF) network

The activator protein-1 (AP-1) family network was obtained by subsetting the APID protein-protein interaction database^87^ down to just the significant AP1-family TFs. Edges between nodes were included if they were supported by at least four experiments. The nodes were color-coded using the signs of the corresponding coefficient of *x_i_*. The network was drawn using Cytoscape^85^.

#### Longitudinal analysis of gene promoter accessibility

We collected promoter tiles from the TSAM of CD16 monocytes in the COVID19L dataset and modeled their accessibility using either GLMMs or ZI-GLMMs with the glmmTMB package^30^. More specifically, for promoters with zeroes, we applied the ZI-GLMM modeling as follows

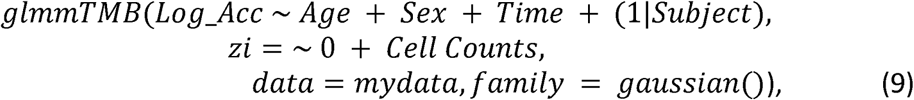

where *Log_Acc* was short for 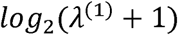, *Time* was days PSO, (1|*Subject*) indicated that random intercepts were used for individual participants, and zero-inflation was modeled as a function of the total cell counts in individual samples with no intercept. For promoters without zeroes, we applied the GLMM modeling as follows

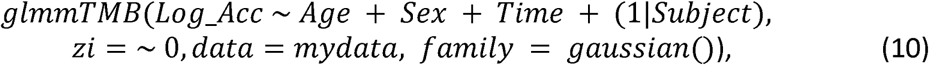

where the ZI component was omitted. The *P* values associated with *Time* were extracted and converted to FDRs to control multiple testing. Promoters with a FDR < 0.1 were considered as significantly changing in time. For promoters attributed to multiple genes, all genes were included for pathway enrichment and downstream analyses.

### Transcription Factor and Gene Promoter Associations

#### Linking Transcription Factor to Gene Promoters

We evaluated whether motif z-scores were statistically associated with gene promoter accessibility via ZI-GLMMs as follows

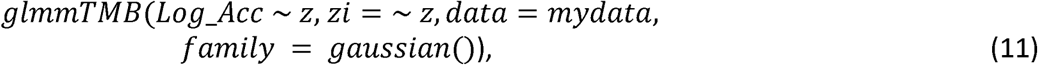

where *Log_Acc* was short for 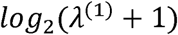 All pairs of significantly changing TFs and significantly changing gene promoters were evaluated. We considered a TF and a gene promoter to be associated if the continuous coefficient of *z* was statistically significant (*P* < 0.05) without adjusting for multiple testing.

#### Linking Transcription Factor to Innate Immune Pathways

Using the TF-gene promoter associations, we calculated the percentage of significant genes in an innate immune pathway were associated with a TF. For visualization purposes, the network only displayed an edge between a TF and a pathway if more than 33% of significant genes in that pathway were associated with the TF.

### Data Availability

The HealthyDonor (GSE190992) and COVID19 (GSE173590) scATAC-seq datasets can be downloaded from the Gene Expression Omnibus (GEO) database under accession numbers GSE190992 and GSE173590, respectively. The corresponding raw data are available via authorized access at dbGaP under accession number phs003203.v1.p1 and phs002576.v1.p1, respectively. The Hematopoiesis dataset was downloaded from (https://www.dropbox.com/s/sijf2votfej629t/Save-Large-Heme-ArchRProject.tar.gz. (Note to Reviewers: The COVID19 dataset will be released to the public prior to the publication of our first manuscript.)

### Code Availability

MOCHA is a freely available R package in CRAN that can be easily downloaded using R or RStudio (https://cran.rstudio.com/web/packages/MOCHA/index.html). All code used to generate figures in this manuscript are available in: https://github.com/aifimmunology/MOCHA_Manuscript(Note to Reviewers: We are currently including this code as a zip file and we will publish all analysis code in Zenodo or equivalent once the content of this manuscript is finalized.)

## Supplementary Figure Legends

**Supplementary Fig. 1.**
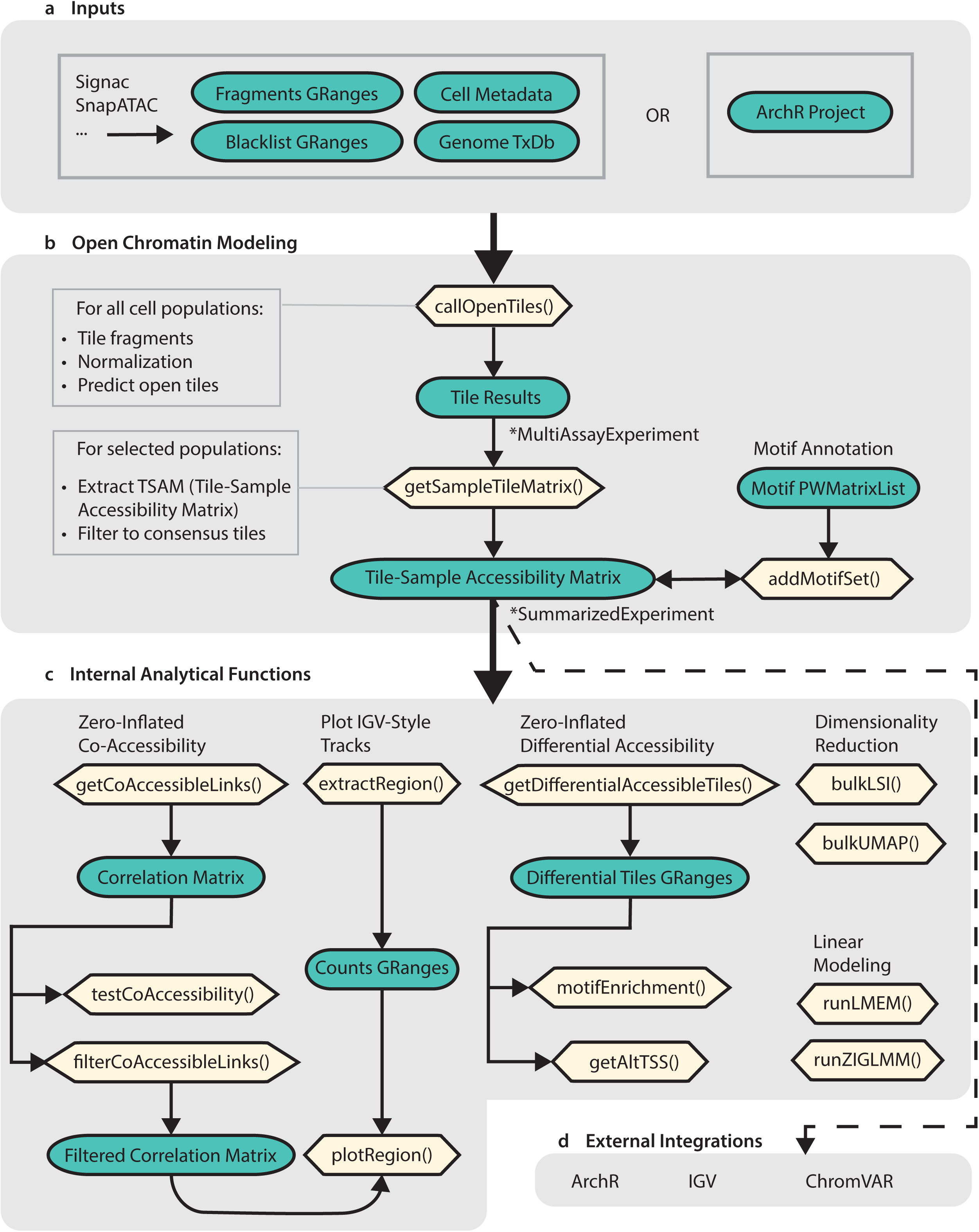
MOCHA’s technical workflow schematic. Schematic of the MOCHA R package workflow functions (in yellow) and objects (green). **a,** MOCHA takes inputs from an ArchR project or collections of input files from ATAC-seq analysis software. **b,** Core functions of MOCHA and result objects. **c,** Downstream analyses supported by MOCHA with functions. **d,** MOCHA enables additional downstream analyses available in external software.

**Supplementary Fig. 2.**
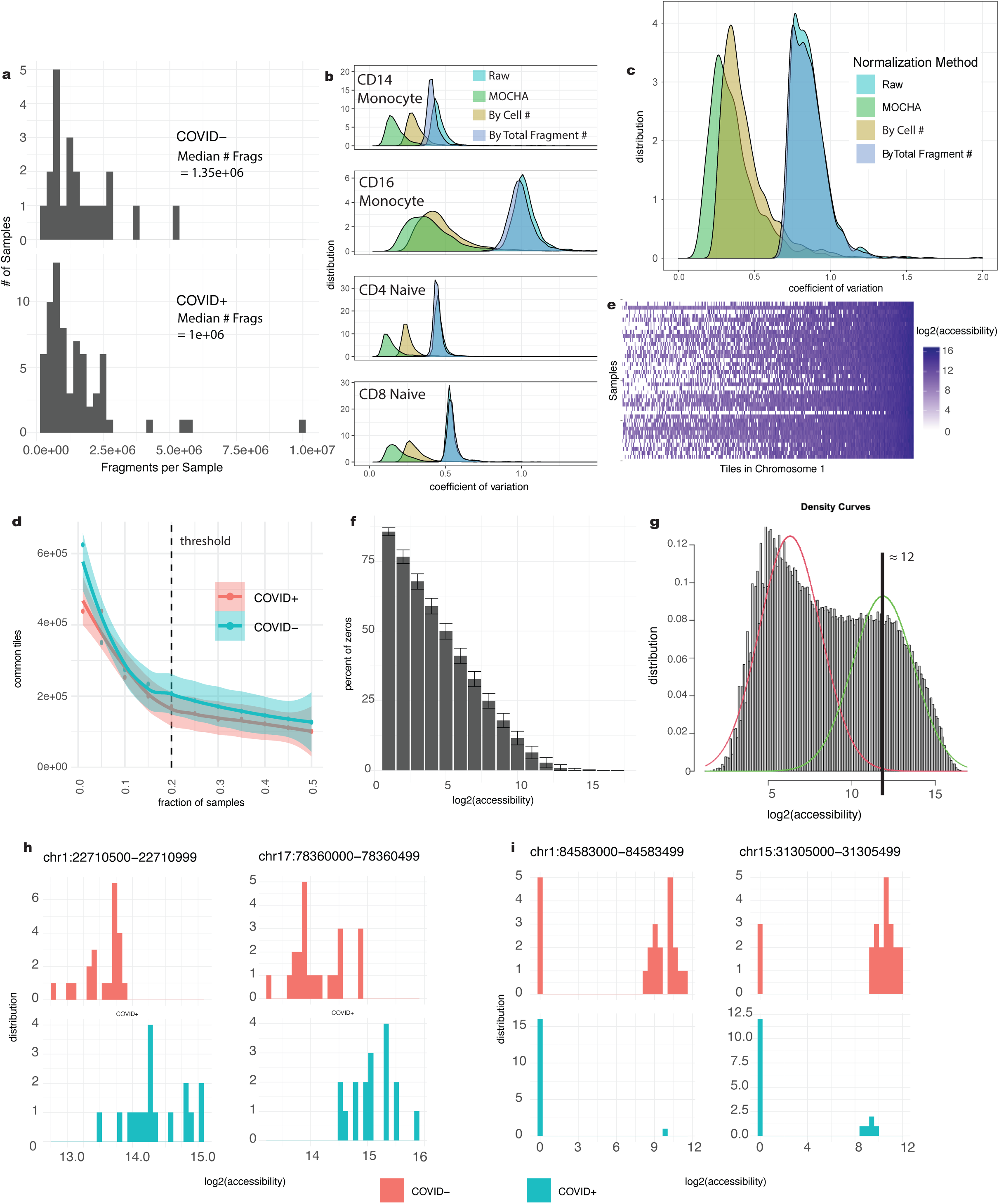
Technical details for developing MOCHA’s analytical modules. **a,** Large difference in sequencing depth per sample was observed in the COVID19 dataset (n=91). The distribution of the number of fragments per sample in the CD16 monocytes are shown separately for COVID+ (n=69) and COVID- samples (n=22). **b-c,** Distributions of coefficients of variation (CVs) of the pseudo-bulk fragment counts at 2,230 cell-type invariant CCCTC-binding factor (CTCF) sites in the COVID19 dataset (n=91) before (Raw) or after normalization by the total number of fragments per cell type per sample (MOCHA), the total number of cells per sample (By Total Cell #), or the total number of fragments per sample (By Total Fragment #). Pseudo-bulk fragment counts were calculated for individual tiles per cell type and sample. For each CTCF site, CV was calculated (**b**) within each cell type across all samples (n=91) or (**c**) across cell types (n=25) within each sample**. d-i,** Based on data in the COVID19X dataset (n=39). **d,** Number of tiles in CD16 monocytes that were commonly open to at least a targeted fraction of samples. Data of COVID+ samples (n=17) and data of COVID- samples (n=22) were analyzed separately. The smooth curves and the shaded bands are the Loess fitting curves and the corresponding 95% confidence intervals. The vertical dashed line indicates the fraction threshold (20%) used. **e,** Heatmap of pseudo-bulk accessibility in the tile-sample accessibility matrix (TSAM) of CD16 monocytes with tiles in Chromosome 1 only. Tiles were sorted by their percentage of zeros across samples. **f,** Histogram of percentage of zeros across samples as a function of tile log2(accessibility) value. The bar represents the mean value while the error bar represents the corresponding standard deviation. **g,** The distribution of accessibilities of all tiles revealing a bimodal distribution. Accessibility threshold was set near the higher mode in order to increase power and avoid testing highly noisy regions in differential accessibility analysis. **h-i,** Exemplar histograms showing differences in accessibility between COVID+ and COVID- samples that arose from either difference in non-zero accessibilities without significant difference in the proportion of zeros (**h**), or difference in the proportion of zeros without significant difference in non-zero accessibilities (**i**). Source data are provided in Source Data Supplementary Fig. 2.

**Supplementary Fig. 3.**
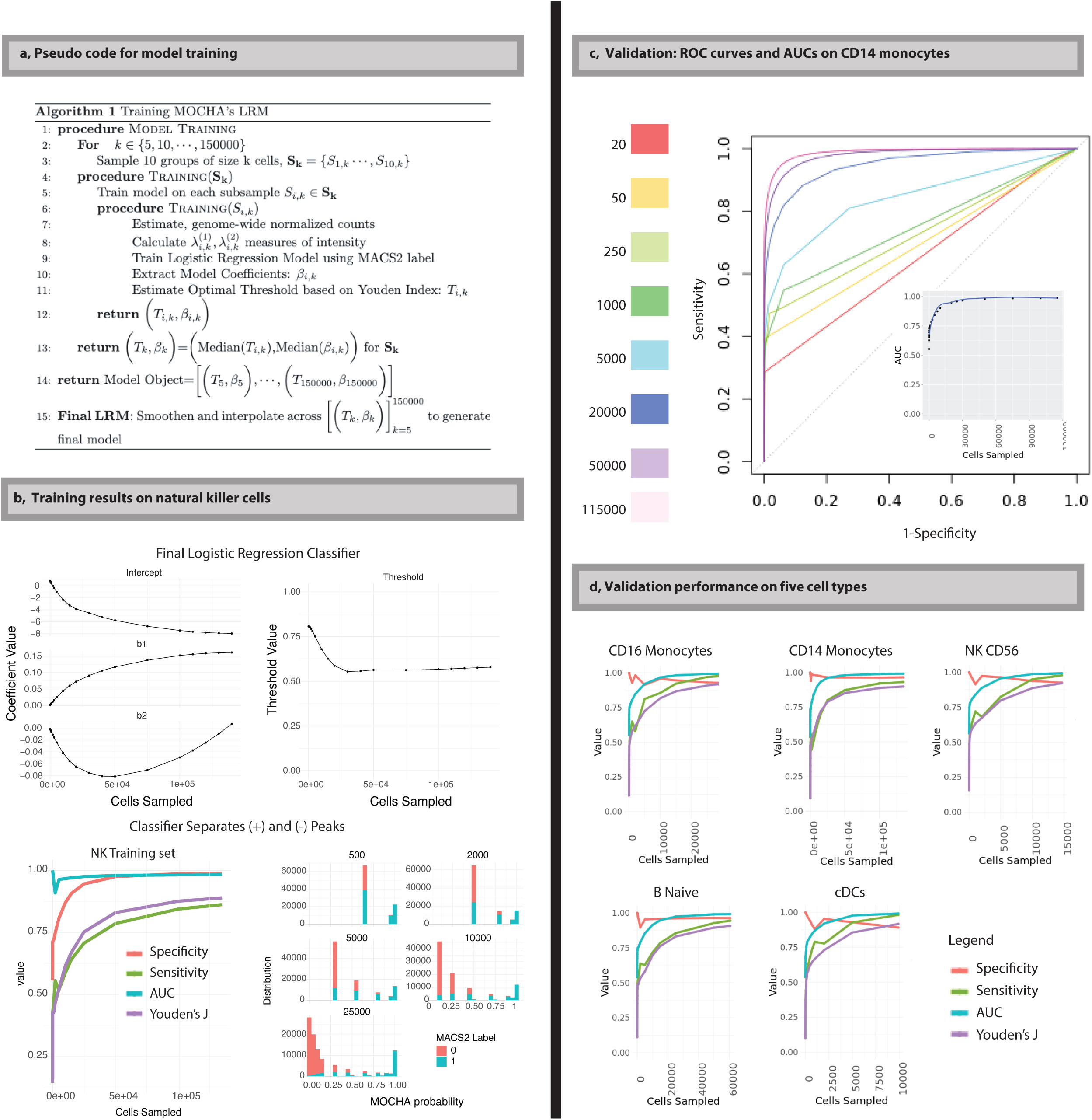
Training and validation of open accessibility models. **a,** Pseudo code for training logistic regression models (LRMs) for open tiles. **b,** Training results on natural killer (NK) cells (n=179,836). Top: Coefficients of the LRMs and the corresponding threshold as a function of sampled cell count. Bottom left: Specificity, sensitivity, area under the receiver operating characteristic (ROC) curve (AUC), and Youden’s J index as a function of sampled cell count. Bottom right: Histograms of the probability scores of open (blue) and closed (red) tiles at cell count 500, 2000, 5000, 10000, and 25000. **c,** ROC curves on the validation data of CD14 monocytes as cell count ranged from 20 to 115,000. Insert: The corresponding AUC as a function of cell count. The Loess fitting curve is plotted in blue. **d,** Validation performance on specificity, sensitivity, AUC, and Youden’s J index as a function of sampled cell count for five representative cell types. cDCs: classical dendritic cells. Data in the COVID19 dataset (n=91 samples) was used for the training and validation of the LRMs. Source data are provided in Source Data Supplementary Fig. 3.

**Supplementary Fig. 4.**
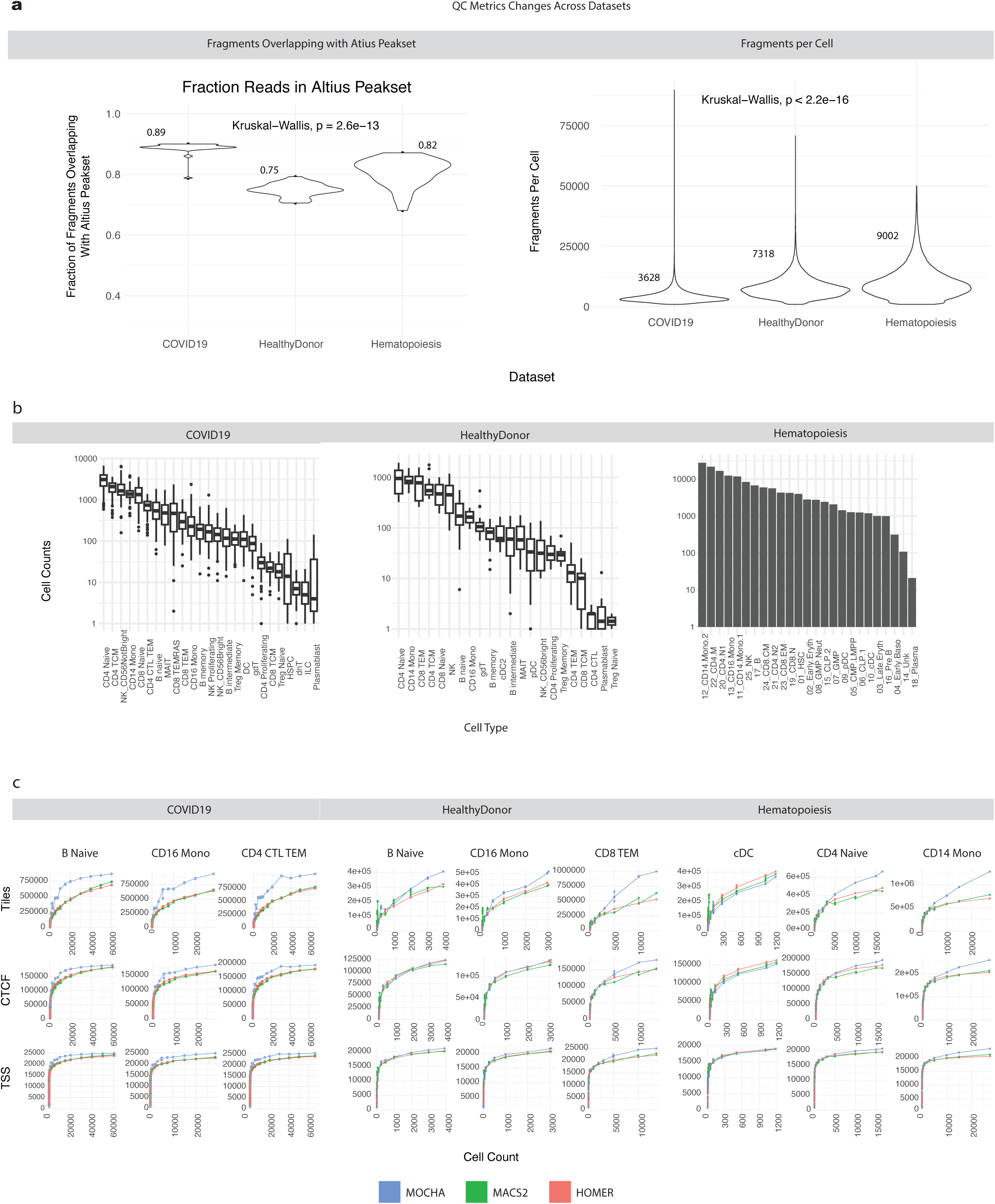
Dataset characteristics and benchmarking on open chromatin identification during downsampling. **a,** Quality control (QC) metrics across three datasets, as measured by the percentage of fragments that overlap with the Altius peakset (left) and the number of fragments per individual cell (right). The corresponding median values are indicated for each dataset. The three datasets were significantly different on these two QC metrics (*P* = 2.6×10^-13^ and *P* < 2.2×10^-16^, respectively; Kruskal-Wallis test). **b,** The number of cells per cell type across the three datasets. Each boxplot displays the median (centerline), the first and third quartiles (the lower and upper bound of the box), and the 1.5x interquartile range (whiskers) of the data. **c**, Head-to-head comparison between MOCHA, MACS2, and HOMER on numbers of detected tiles (top), tiles overlapping with CTCF sites (middle), and overlapping with TSSs (bottom) as functions of sampled cell count in three representative cell types from each of the three datasets. The three datasets are COVID19 (n=91, left), HealthyDonor (n=18, middle), and Hematopoiesis (treated as n=1 sample, right). CTCF: CCCTC-binding factor; TSS: transcription starting site. Source data are provided in Source Data Supplementary Fig. 4-1 and 4-2.

**Supplementary Fig. 5.**
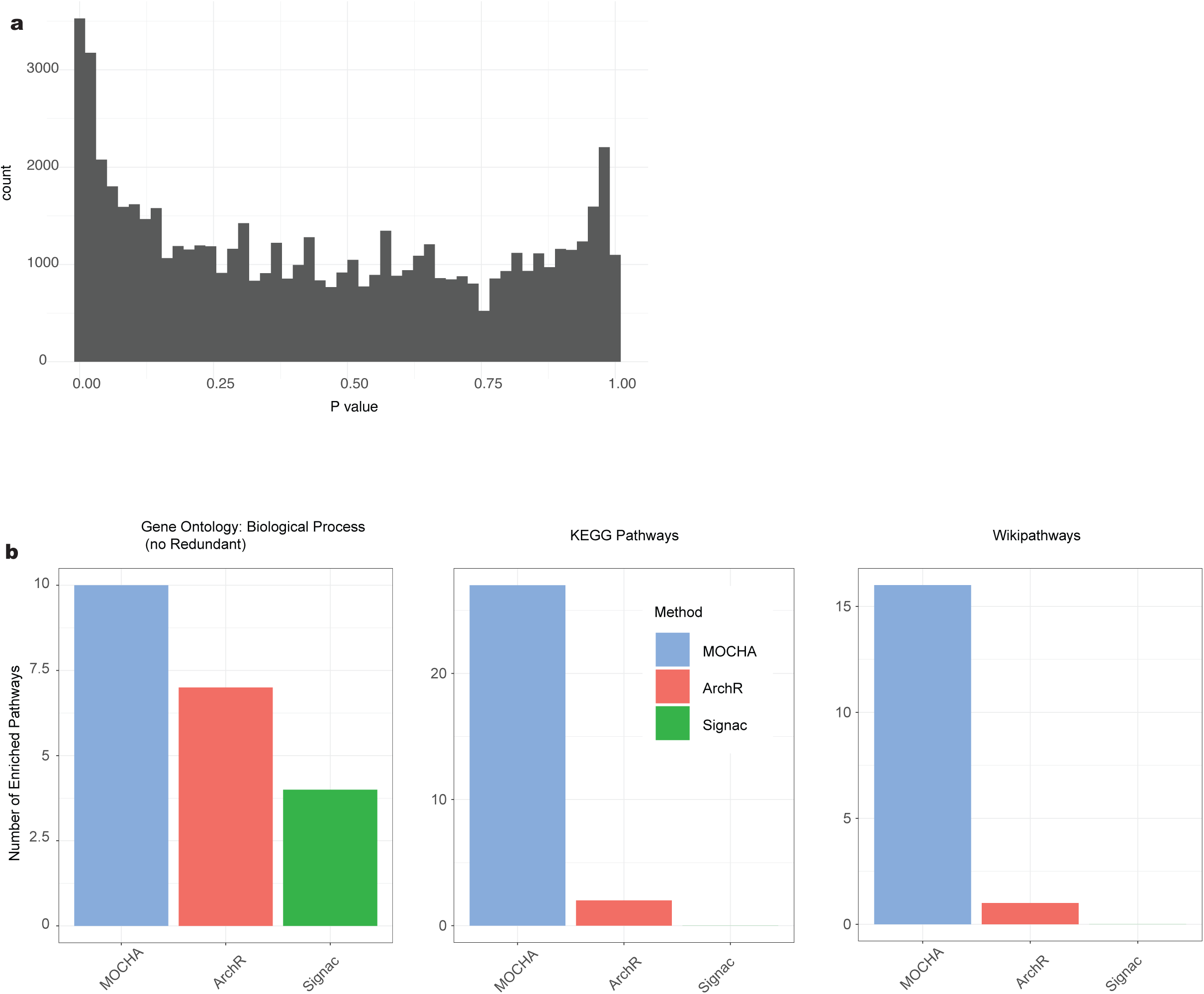
Additional information on differential accessibility analysis. **a,** Histogram of *P* values evaluated by MOCHA on filtered open tiles. Only *P* ≤ 0.95 values were used for estimating false discovery rate (FDR). **b,** Pathway enrichment analysis on genes having differential accessibility tiles (DATs) in their promoter regions using the Gene Ontology (left), KEGG (middle), or Wikipathway (right) database. The DATs were identified by MOCHA, ArchR, or Signac. Source data are provided in Source Data Fig. 3-S5.

**Supplementary Fig. 6.**
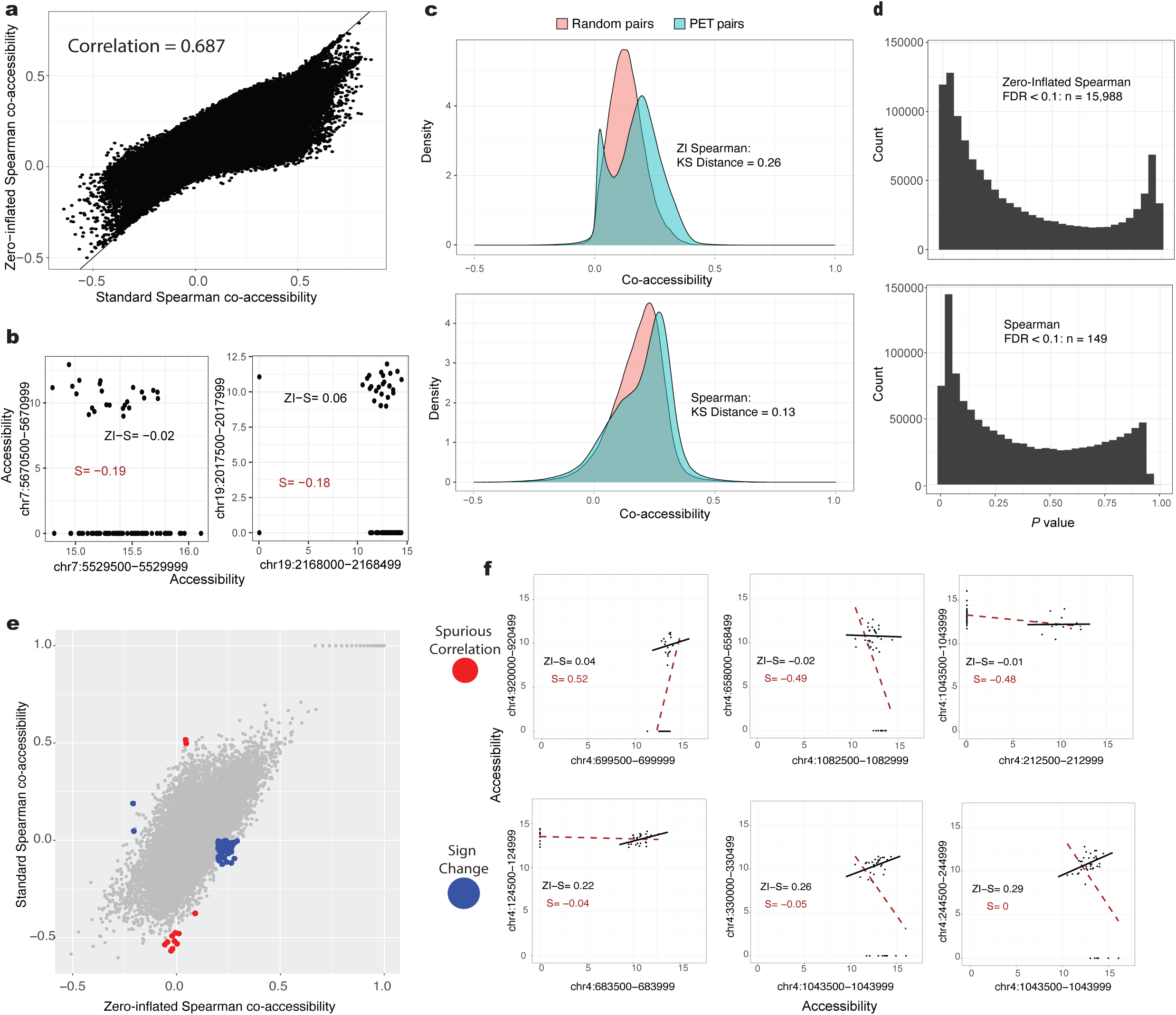
Comparing standard Spearman and zero-inflated Spearman in evaluating co-accessibility. **a,** A direct comparison between standard Spearman (S) and zero-inflated Spearman (ZI-S) in evaluating inter-cell type co-accessibility between tile pairs linked to promoter-enhancer interactions in naive CD8 and CD4 T cells^43^. Co-accessibilities between individual promoter-enhancer tile (PET) pairs (n=1.2 million) were conducted across 17 cell types in the full COVID19 dataset (n=91 samples). **b,** Examples of divergence between the zero-inflated Spearman and the standard Spearman co-accessibilities on two PET pairs. Each dot represents one cell type and sample combination (n=1547 data points). **c,** Distributions of co-accessibilities between PET pairs and randomly selected tile pairs (n=100k) as evaluated by ZI-Spearman (top) or Spearman (bottom). The Kolmogorov-Smirnov (KS) distance was used to quantify the separation between the distributions of PET pairs and random pairs. **d,** Histogram of empirical *P* values of PET pairs that were calculated based on the percentile of random pairs, using either ZI-Spearman (top) or Spearman (bottom). False discovery rate (FDR) estimation was conducted for each *P* value set. The number of PET pairs with FDR < 0.1 is shown. **e,** Comparison of inter-sample co-accessibility within CD16 monocytes in the COVID19X dataset (n=39 samples) as evaluated by standard Spearman or ZI-Spearman. All possible pairs of tiles within the first million base pairs of Chromosome 4 were evaluated for an illustrative purpose. **f,** Examples of spurious correlations (top) and sign changes (bottom) generated by standard Spearman correlations on ZI data, as compared to results from the ZI-Spearman. Source data are provided in Source Data Supplementary Fig. 6-1 and 6-2.

**Supplementary Fig. 7.**
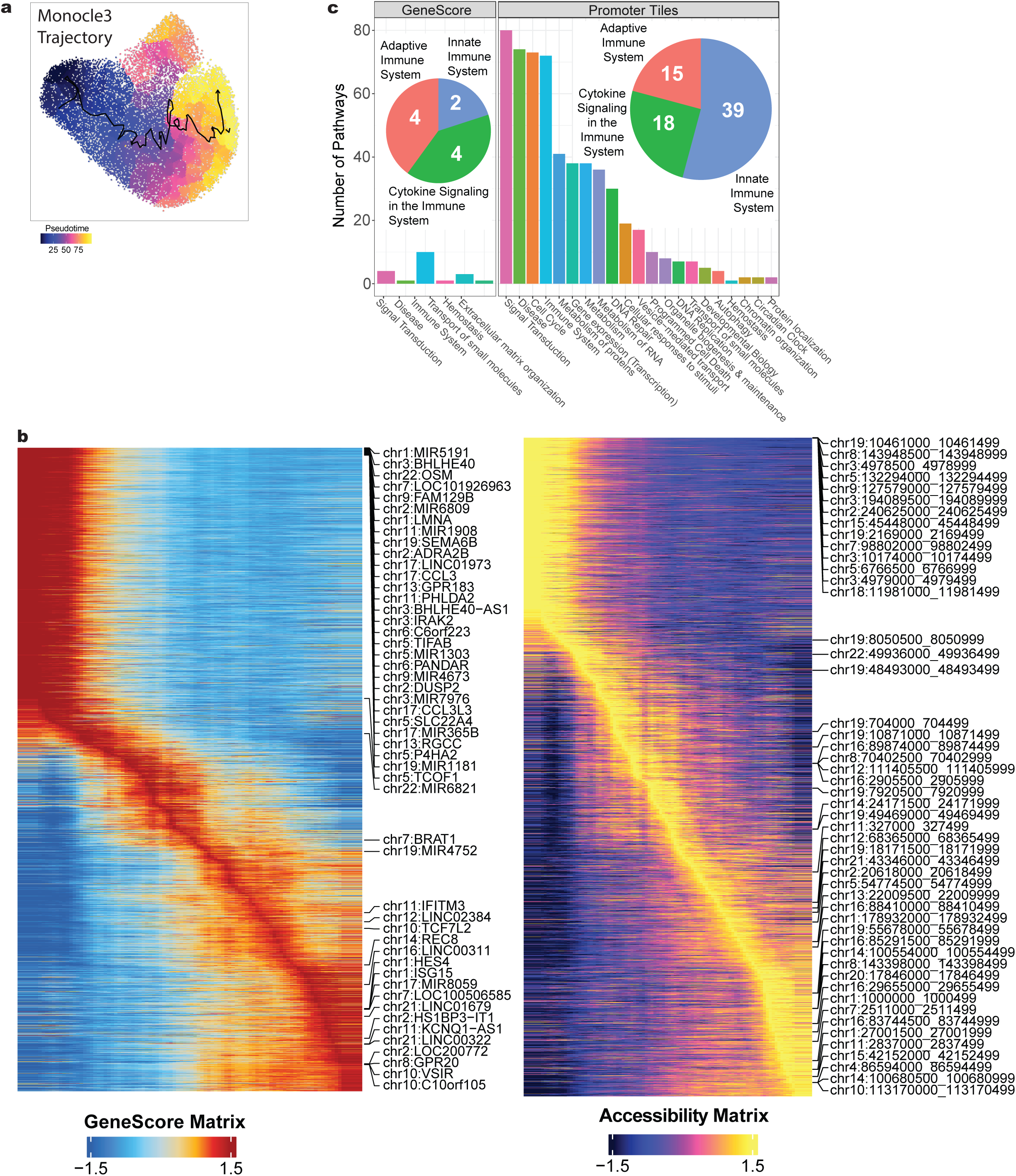
Benchmarking pseudotime trajectory analysis between MOCHA’s tiles and ArchR’s genescores. **a,** Monocle3 trajectory constructed from CD16 monocytes belonging to samples in the order of early infection (1–15 days PSO, n=21), late infection (16–30 days PSO, n=13), recovery (>30 days PSO, n=35), and uninfected (n=22) in the COVID19 dataset. The trajectory is overlaid on the corresponding single-cell UMAP. **b,** Pseudotime heatmaps of ArchR’s genescores (left) and MOCHA’s accessible tiles (right) that were generated using ArchR standard settings. The top 50 genes or tiles, respectively, are labeled. **c**, Significant Reactome pathways (FDR < 0.05) enriched with genes having highly variable genescores or promoter accessibility changes along the pseudotime trajectory. The variability threshold was set using ArchR’s standard threshold (varCutOff = 0.9). The pathways were aggregated into upper-level pathway annotations using Reactome’s database hierarchy. The barplot shows the number of pathways in each category. The pie chart breaks down the immune system pathways by Reactome’s next level categories. PSO: post symptom onset; FDR: false discovery rate. Source data are provided in Source Data Supplementary Fig. 7-8.

**Supplementary Fig. 8.**
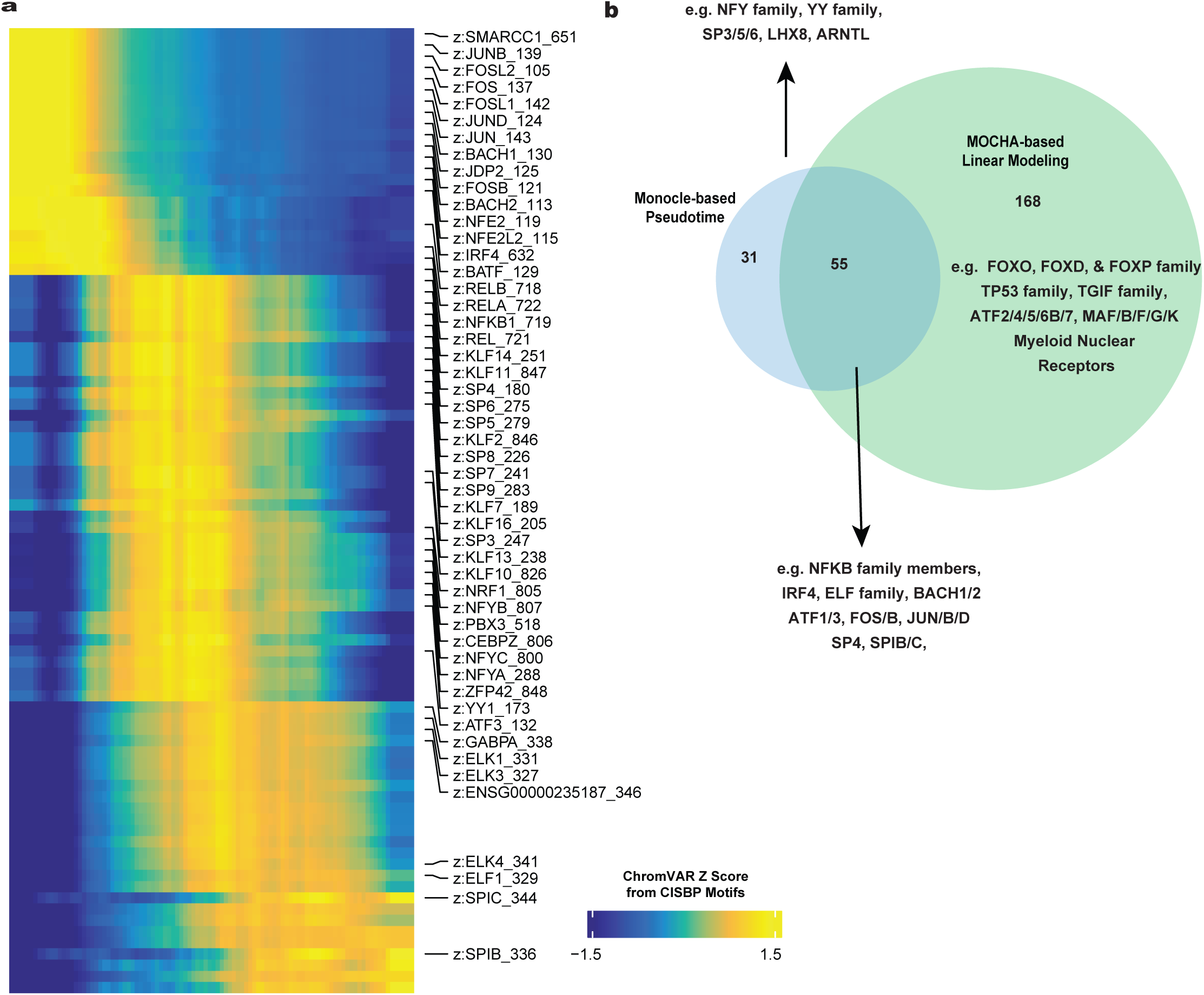
Benchmarking motif usage between single-cell level pseudotime trajectory analysis and pseudo-bulk level real time longitudinal modeling. **a,** Pseudotime heatmap of ChromVAR z-scores along the trajectory constructed on CD16 monocytes in the COVID19 dataset (n=91 samples). ChromVAR z-scores were evaluated based on the CISBP database. The top 50 motifs are labeled. Uninfected samples were excluded from the analysis. **b,** Venn diagram comparing motifs showing significant ChromVAR z-score changes either at single-cell level along the pseudotime trajectory (ArchR standard threshold) or at pseudo-bulk level in real time (days PSO) as modeled by generalized linear mixed models (FDR < 0.1). Motif examples for each subsets are provided. FDR: false discovery rate. PSO: post symptom onset. Source data are provided in Source Data Supplementary Fig. 7-8.

**Supplementary Fig. 9.**
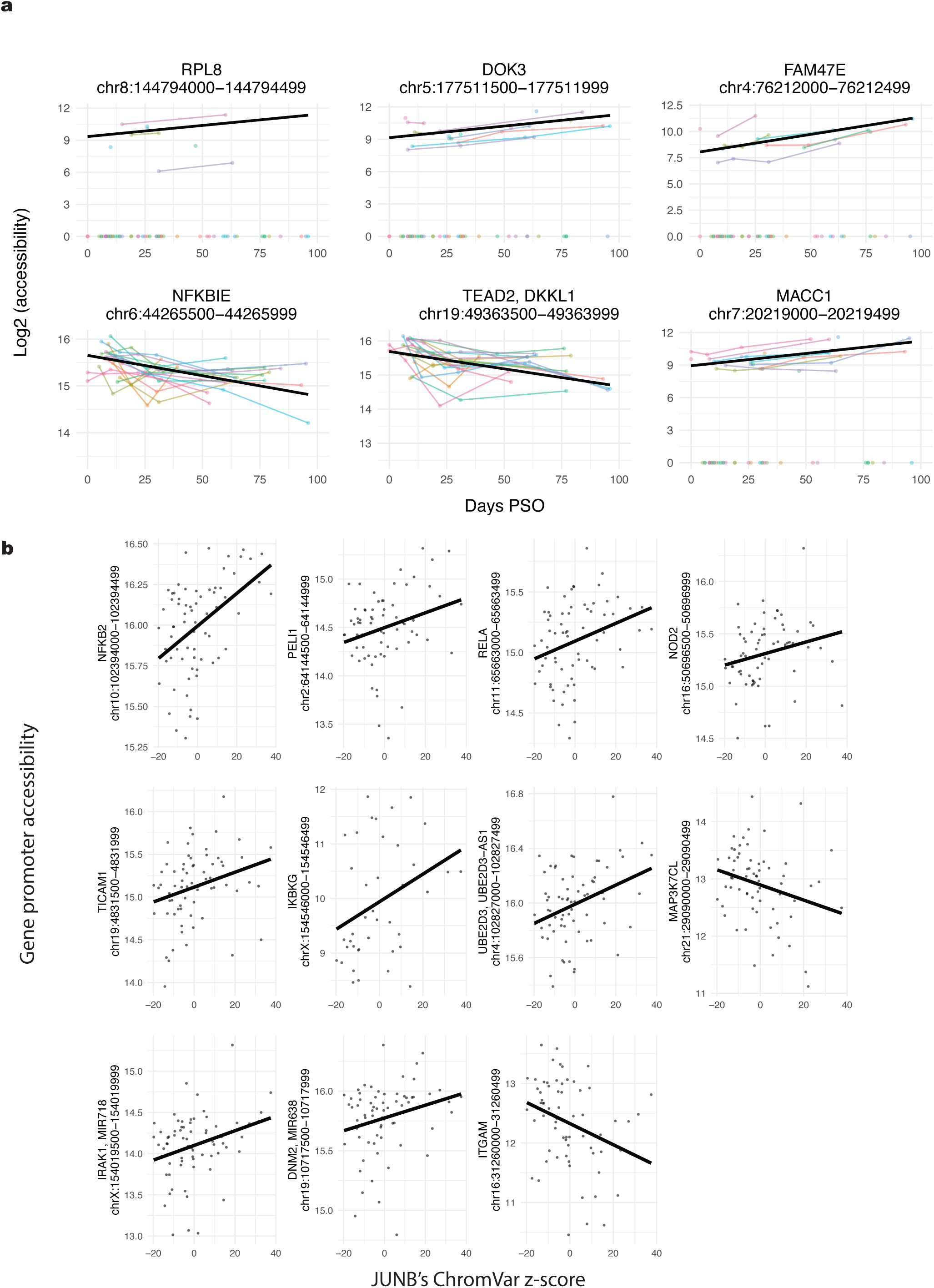
Examples illustrating longitudinal shifts in gene promoter accessibility and motif-promoter associations during COVID-19 recovery. **a,** The top 6 genes showing significant promoter accessibility changes (FDR < 0.1) based on ZI-GLMM. Data of individual participants are shown in thin colored lines. **b,** Scatter plots illustrating examples of significant associations (P < 0.05) between JUNB’s ChromVAR z-score and significantly changing (FDR < 0.1) promoter accessibility of genes within the TLR4 Reactome pathway. **a-b,** Based on the COVID19L dataset (n=69 samples). The thick black line shows the population trend from ZI-GLMM. PSO: post symptom onset; FDR: false discovery rate; ZI-GLMM: zero-inflated generalized linear mixed effects model; TLR4: toll-like receptor 4. Source data are provided in Source Data Supplementary Fig. 9.

## Supplementary Table Legends

Supplementary Table 1. Literature references for all Type II alternatively regulated genes, which are indicated as either altered in COVID-19 or identified as potential therapeutic targets.

## Source Data

**Source Data Fig. 2-1**

Source data for Fig. 2a,b,d-h.

**Source Data Fig. 2-2**

Source data for Fig. 2c (zipped BED files).

**Source Data Fig. 3-S5**

Source data for Fig. 3 and Supplementary Fig. 5.

**Source Data Fig. 4**

Source data for Fig. 4a,d-g.

**Source Data Fig. 5**

Source data for Fig. 5.

**Source Data Supplementary Fig. 2**

Source data for Supplementary Fig. 2.

**Source Data Supplementary Fig. 3**

Source data for Supplementary Fig. 3b-d.

**Source Data Supplementary Fig. 4-1**

Source data for Supplementary Fig. 4a (left pane), b-c.

**Source Data Supplementary Fig. 4-2**

Source data for Supplementary Fig. 4a (right panel).

**Source Data Supplementary Fig. 6-1**

Source data for Supplementary Fig. 6a,c-d.

**Source Data Supplementary Fig. 6-2**

Source data for Supplementary Fig. 6b,e-f.

**Source Data Supplementary Fig. 7-8**

Source data for Supplementary Fig. 7b,c and Supplementary Fig. 8.

**Source Data Supplementary Fig. 9**

Source data for Supplementary Fig. 9.

## Acknowledgements

We thank the study participants for their dedication to this project; the Allen Institute founder, Paul G. Allen, for his vision, encouragement, and support; Allan Jones for his support to our COVID-19 study; Qiuyu Gong for providing code for plotting single-cell density within UMAP space; Ziyuan He, Peiyao Zhang, Marla Glass, and Sandra Munro for review and helpful discussions of the manuscript draft; Leila Shiraiwa and Nina Kondza for laboratory operations support; the Human Immune System Explorer (HISE) software development team at the Allen Institute for Immunology for their support and dedication. This paper and the research behind it would not have been possible without the collaborative computational data analysis environment provided by HISE. The research reported in this publication was supported in part by COVID supplements from the National Institute of Allergy and Infectious Diseases and the Office of the Director of the National Institutes of Health under award numbers UM1AI068618-14S1 and UM1AI069481-14S1 (MJM). This work was also supported by Paul G. Allen Family Foundation Award #12931 (MJM); Seattle COVID-19 Cohort Study (Fred Hutchinson Cancer Research Center, MJM); and the Joel D. Meyers Endowed Chair (MJM). The content is solely the responsibility of the authors and does not necessarily represent the official views of the funders.

## Author Contributions

S.R.Z, M.P.P, and X.-j.L. conceived the study and jointly designed the development, benchmarking, and analysis. M.J.M. and T.F.B. acquired the funding. T.R.T, P.J.S., X.-j.L., M.J.M., T.F.B. designed the COVID-19 study. J.L.C. and M.J.M. enrolled participants and collected their samples. M.W., J.R., P.J.S. processed the samples and performed the scATAC-seq experiment.

S.R.Z., M.P.P., I.M., and L.O. developed the software package. I.M. led the CRAN submission. S.R.Z, M.P.P, and X.-j.L. analyzed the data. X.-j.L. supervised and directed the project. S.R.Z, M.P.P, and X.-j.L. wrote the manuscript. All authors provided edits and comments to the manuscript.

## Competing interests

The authors declare no competing interests.

## Notes

### Competing Interest Statement

The authors have declared no competing interest.

